# A histidine switch controls the pH-responsive self-assembly of a helical protein filament

**DOI:** 10.64898/2026.06.13.732084

**Authors:** Swasti Rawal, Stefan Bohn, Maria Bacia-Verloop, Benjamin Bourgeois, Ðesika Kolarić, Dagmar Kolb, Tea Pavkov-Keller, Iva Pritišanac, Tobias Madl, Ambroise Desfosses, T. Reid Alderson

## Abstract

Self-assembling helical protein filaments underlie diverse biological processes, from signaling pathways to cell motility. Encoding tunable self-assembly into the sequences of filamentous proteins remains a major challenge. Here, we discovered that the caspase-9 CARD can natively self-assemble into helical filaments in a pH-regulated manner. We defined the determinants of filament assembly using an integrative structural, biophysical, and computational approach. Using NMR spectroscopy, we found that the protonation of a single histidine residue near an N-terminal helix dipole, H38, regulates the pH-dependent self-assembly process. Charge-altering mutations at this site tune thermodynamic stability and filament self-assembly across solution and pH conditions. We solved 3.3- and 3.5-Å cryo-EM structures of the wild-type and H38R filaments, respectively, which show H38 positioned directly at a filament interface. Molecular dynamics simulations show that H38 functions as a molecular switch, whereby protonation rotates its positively charged side-chain toward solvent and away from the partial-positive charge at the N-terminal helix dipole. This reflects a fine balance between stabilizing intermolecular association and a destabilizing intramolecular electrostatic clash at the helix dipole. More broadly, across 350 helix-containing protein domains we identified electrostatic contributions to protein stability near helix dipoles by integrating AlphaFold2 predictions with deep mutational scanning data. Together, our results identify a native, pH-sensitive histidine switch that regulates a self-assembling helical protein filament. Our results establish a mechanism by which charge-altering mutations near helical N-termini can be engineered to control side-chain rotamers, protein stability, and self-assembly.

## Introduction

The dynamic assembly of protein complexes plays a central role in transmitting biological information^1^. Protein self-assembly, including the formation of higher-order signaling complexes or signalosomes^2–4^, is a widespread mechanism that initiates cell-signaling pathways and propagates downstream signals.^1,5,6^ By increasing local concentrations and enabling multivalent interactions, protein oligomerization can dramatically enhance the efficiency of molecular recognition and enzymatic activity^7^. An extreme case of self-assembly is the polymerization of protein filaments^8^, which can be pathological in diseases such as sickle cell anemia^9^ and neurodegeneration^10^. However, functional protein filaments are widespread throughout biology and regulate metabolic enzyme activity^11–13^ and signaling processes involving cytoskeletal organization^14,15^, flagellar motility^16^, autophagy^17^, prokaryotic antiviral defense^18–20^, and the nutrient starvation response^21,22^.

Synthetic or designed protein filaments, often inspired by natural processes such as filamentous spider silk assembly^23^, find broad use as nanomaterials^24,25^ and in biotechnology for metabolic engineering and cellular organization^26,27^. However, encoding environmentally tunable self-assembly into the sequences and structures of protein filaments remains challenging^24,25,28,29^. Recently, a *de novo* design strategy based on networks of buried histidine residues was used to tune the pH-responsive assembly of both homo-oligomers and helical protein filaments^25,30^. This approach, however, requires extensive computation and *in-vitro* screening of designed variants. In naturally occurring proteins, only a handful of pH-gated protein filaments have been described^31–33^ and are largely restricted to metabolic enzymes under conditions of cell starvation^31^. Moreover, the mere presence of a titratable histidine residue at a filament interface does not necessarily confer pH sensitivity^32^, suggesting that pH-regulated filament assembly may preferentially emerge in proteins that both contain histidine and suitably charged residues at the filament interface.

Death domain folds (DDFs) are small protein domains that contain a high content of charged residues with generally polarized surface. Numerous proteins in the DDF superfamily have been reported to self-assemble into helical filaments^34–43^. DDF filament formation by cysteine-aspartic proteases^44,45^ (caspases), including caspase-1 and caspase-8, triggers apoptotic and inflammatory signaling via proximity-induced dimerization and activation of the covalently attached protease domains^35,46,47^. DDFs fold into conserved six-helix bundles and mediate protein-protein interactions in apoptotic and inflammatory signaling pathways^48–50^. Among the different DDFs, the caspase activation and recruitment domain (CARD) sub-family of DDFs has been extensively studied, with locally asymmetric binding interfaces among CARD complexes that can lead to complex modes of homo- and hetero-oligomerization, such as polymerization into helical filaments^34–43^ and the formation of biomolecular condensates via liquid-liquid phase separation^51^. However, the molecular mechanisms of CARD and DDF self-assembly, particularly the events that trigger filament formation and how this is regulated by environmental cues, remain unclear.

Here, we discovered that the caspase-9 CARD (C9^CARD^) self-assembles into helical protein filaments in a pH-responsive manner controlled by a single histidine residue, H38. We solved a 3.3-Å cryo-EM structure of the wild-type C9^CARD^ filament, in which H38 is situated at a filament interface. Charge-altering mutations at this position (H38R, H38D, H38N) dramatically modulate the propensity to self-assemble into filaments. We subsequently determined a 3.5-Å cryo-EM structure of the H38R variant of C9^CARD^, which serves as a mimic of the histidine-protonated form. To better understand the impact of H38 protonation or H38R mutation, we performed all-atom molecular dynamics (MD) simulations of the C9^CARD^ and observed reorganization of the side-chain rotameric states at this position, with H38 protonation and the H38R mutation favoring orientations away from the helix dipole toward solvent. We leveraged AlphaFold2 to analyze helix residue frequencies across all human protein domains, and with deep mutational scanning (DMS) data from 350 protein domains, we found that introducing a positive charge near helical N-termini is broadly destabilizing. Our results collectively show that titratable His residues near the N-terminal helix dipole, such as H38 in the C9^CARD^, may serve as exquisite pH sensors that can regulate protein stability, side-chain rotameric states, and self-assembly.

## Results

DDFs often self-assemble into helical filaments through highly charged, polarized interfaces ^34–43^. We asked whether titratable histidine residues could function as pH sensors that gate helical filament formation in DDFs, thereby tuning self-assembly near physiological pH. The DDF-containing caspase-1 forms filaments via its CARD domain^52,53^ (C1^CARD^) (**Figure 1A**), which comprises 30% charged residues overall but is devoid of titratable His residues. We therefore focused on its paralog caspase-9 (C9^CARD^) (**Figure 1B**), which forms stoichiometric complexes with Apaf-1 in the apoptosome (**Supplementary Appendix**) and contains only a single His residue. However, the C9^CARD^ has not been reported to polymerize, despite extensive biochemical and structural studies^54–58,58–72,72,73,73–77^. We therefore tested whether the C9^CARD^ may self-assemble like its paralog, and if its lone His residue may function as an endogenous pH sensor for self-assembly.

**Figure 1.**
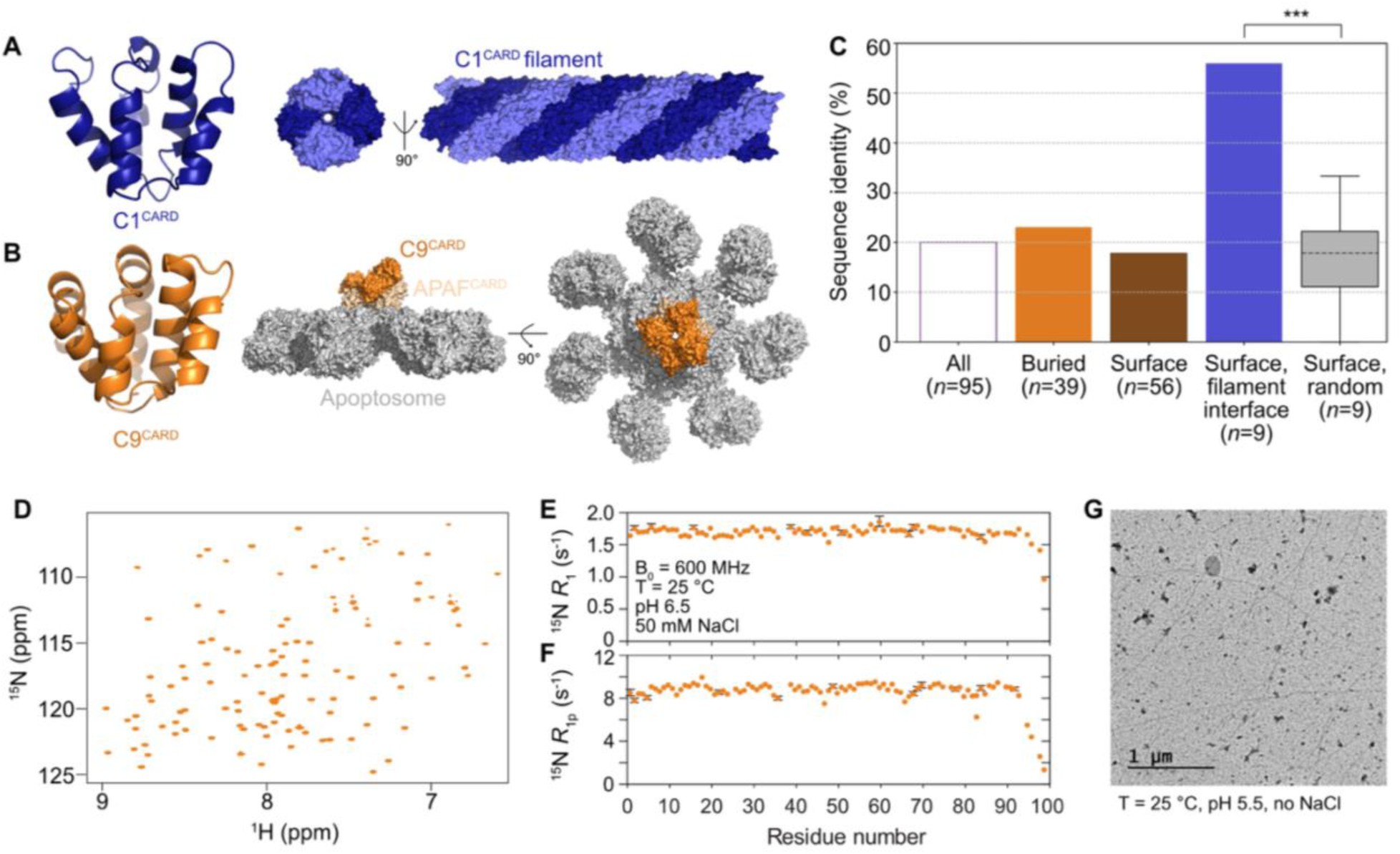
The C9^CARD^ self-assembles into filaments. (**A**) Structure of the C1^CARD^ depicted as a monomer (PDB: 5fna, chain A) and in the filamentous state. (**B**) Structure of the C9^CARD^ (PDB: 4rhw, chain E) and when bound to the apoptosome platform (PDB: 5wve). The C9^CARD^ is colored orange, the APAF^CARD^ in light orange, and the other components and domains of the apoptosome in grey. (**C**) Pairwise sequence identity between C1^CARD^ and C9^CARD^ computed for all residues in the alignment (white), only buried residues (orange, rSASA < 25%), only surface residues (brown, rSASA ≥ 25%), or only surface residues involved in the largest C1^CARD^ filament interface (light blue). The distribution of sequence identities among randomly sampled surface residues (grey, 10,000 simulations) is shown in a box plot. The number of residues in each category is listed. *** indicates a p value of 1e-4. (**D**) 2D ^1^H-^15^N HSQC spectrum of uniformly ^15^N-labeled C9^CARD^ at 1 GHz in 25 mM HEPES, 50 mM NaCl, 5 mM TCEP, 0.5 mM EDTA at pH 6.5 and 25 °C. (**E**) ^15^N *R*_1_ and (**F**) ^15^N *R*_1ρ_ relaxation rates in C9^CARD^ recorded at 25 °C and a static magnetic field strength of 14.1 T (600 MHz). The ^15^N spin-lock field strength used for the *R*_1ρ_ experiment was 2 kHz. Errors represent one standard deviation from the mean and were derived from analysis of planes with duplicated delay times in the relaxation datasets. (**G**) Representative negative-stain EM image of C9^CARD^ filaments that were obtained from a sample in the absence of salt at pH 5.5 where the maximum solubility of C9^CARD^ is 150 μM.

Alignment of the C9^CARD^ and C1^CARD^ structures yields a backbone root-mean-square deviation of 2.0 Å, which indicates highly similar folds (**Figure 1A**, **Figure 1B**). However, a global pairwise sequence alignment of the C9^CARD^ and C1^CARD^ reveals only 20% sequence identity (**Figure 1C**). This corresponds to 77 amino-acid differences among the 96 aligned positions. Moreover, the pairwise sequence identity does not appreciably differ when considering only buried residues or only solvent-exposed residues (**Figure 1C**), indicating that the sequence differences are widespread throughout the domains. This contrasts with sequence comparisons between C1 (or C9) and its orthologs, where buried residues are far more likely to be conserved (**Supplementary Figure 1**). Thus, the sequences of the paralogous C1^CARD^ and C9^CARD^ domains have highly diverged despite retaining structural similarity.

Detailed inspection of the C1^CARD^ and C9^CARD^ surfaces reveals more nuanced insight into their evolutionary relationships. Nine surface residues in C1^CARD^ comprise the largest filament interface and therefore contribute most to its stability^35^ (PDB: 5fna; R10, K11, R15, N23, D27, E28, Q31, D52, R55). We observed that the C9^CARD^ and C1^CARD^ exhibit 56% identity at these nine positions (**Figure 1C**). To determine a baseline expectation value, we iteratively sampled surface residues (*n* = 9) at random from the C1^CARD^ and quantified the pairwise sequence identity to C9^CARD^ at these sites, yielding a mean sequence identity of 18 ± 12%. When compared to the observed 56% identity above, this random sampling approach corresponds to a Z-score of 3.2 and an empirical p value of 1 x 10^-4^, indicating that sequence conservation at the C1^CARD^ filament interface is not due to chance alone. This observation is surprising because surface residues typically evolve at a 4-fold faster rate than buried residues^78,79^. Conserved surface residues can signify shared and functionally relevant protein-protein interaction sites; however, the C1^CARD^ and C9^CARD^ do not share any known DDF interactors^80^. An alternative explanation is that the conserved surface residues found in C9^CARD^ are preserved to maintain a homo-oligomeric interface, e.g. the type-1 interface of a CARD filament. Our sequence and structural analyses led us to hypothesize that C9^CARD^ may self-assemble in a similar manner to other DDFs.

### C9^CARD^ self-assembles into filaments

To examine the self-assembly propensity of C9^CARD^ in more detail, we expressed and purified recombinant, ^15^N-labeled labeled C9^CARD^ for NMR spectroscopy (**Figure 1D**). The two-dimensional ^1^H-^15^N heteronuclear single-quantum coherence (HSQC) spectrum of the ^15^N-labeled C9^CARD^ at pH 6.5 and room temperature confirmed that our construct is folded under these conditions with the expected number of backbone amide signals. We also observed an additional 10 signals from aliased Arg Nε-Hε groups, which exhibit ^15^N chemical shifts near 85 ppm, as confirmed by increasing the ^15^N spectral width (**Supplementary Figure 2**). Next, we recorded triple-resonance NMR data on a ^13^C,^15^N-labeled sample and assigned the backbone and Cβ resonances. The secondary ^13^Cα chemical shifts indicate that the six helices are formed in solution (**Supplementary Figure 3**), including a kink in helix-1. Moreover, ^15^N spin relaxation experiments confirm that the C9^CARD^ is monomeric in solution under these conditions with a rotational correlation time of approximately 6.5 ns (**Figure 1E**, **Figure 1F**). This falls between an empirical value of 7 ns for a protein of this molecular mass^81^ (from a standard curve obtained 5 °C lower than our conditions), as well as the value predicted by HydroNMR^82^ (5.5 ns, **Methods**) using a crystal structure of C9^CARD^ that lacks six residues found in our construct. To test for transient oligomerization that may affect the rotational correlation time, we also measured ^15^N Carr-Purcell-Meiboom-Gill relaxation dispersion experiments; however, we did not detect any millisecond-exchange contributions to the ^15^N transverse relaxation rate (**Supplementary Figure 4**). Thus, the C9^CARD^ remains folded and monomeric under these conditions, which is consistent with previous structural studies.

During the NMR sample preparation, we noticed that C9^CARD^ began to aggregate near *ca.* 0.3 mM at pH 6 in salt-free buffers. This prompted us to use negative-stain (ns) EM to examine the properties of aggregates obtained in salt-free buffer at pH 7 and observed filament-like structures with an average diameter of approximately 10 nm (**Supplementary Figure 5**). Similar filaments were obtained when the C9^CARD^ was prepared in salt-free buffer at acidic pH (**Figure 1G**), although a lower protein concentration was required to induce aggregation. Thus, the C9^CARD^ self-assembles into filaments, which has not been reported previously in various investigations into its structural and biophysical properties^58,72,73,76,83,84^.

Because our experiments used only the isolated CARD domain, we tested whether inclusion of the disordered interdomain linker (residues 100-138) and the protease domain (residues 139-416) impacted filament formation. We purified the C9^CARD+linker^ construct (residues 1-138) and full-length C9 (1-416) harboring an inactivating C287A mutation that prevents auto-proteolysis of the p20-p10 linker^85^. We collected negative-stain EM data to visualize the aggregates formed by these longer constructs, finding that the majority of aggregates were amorphous, but several filaments could be identified (**Supplementary Figure 6**). Thus, we proceeded to characterize the isolated C9^CARD^ filaments henceforth given ease with which filaments assemble.

### Cryo-EM structure of the C9^CARD^ filament

To examine the structural properties of the C9^CARD^ filament in more detail, we determined a high-resolution structure of the filament with cryo-EM (**Figure 2A**). We observed bundle formation in which multiple filaments clustered together with flexible and dynamic morphology (**Supplementary Figure 7**). During cryo-EM data processing, however, we were able to overcome the issue of bundle formation, eventually obtaining a 3.3-Å map (**Figure 2A, Supplementary Figure 7**). Our cryo-EM derived model of the C9^CARD^ subunit is highly similar to the available crystal structures of the C9^CARD^, with backbone root-mean-square deviation values near 0.7 Å and 0.6 Å, respectively, showing that the overall tertiary structure is preserved within the filament. The C9^CARD^ filament displays *C*_2_ symmetry with left-handed, two-start helical assembly consisting of approximately five subunits per turn and a twist angle of -68.0° and an axial rise of 9.3 Å per subunit. The inner and outer diameters of filament are respectively 2.3 and 9.5 nm (**Figure 2A**).

**Figure 2.**
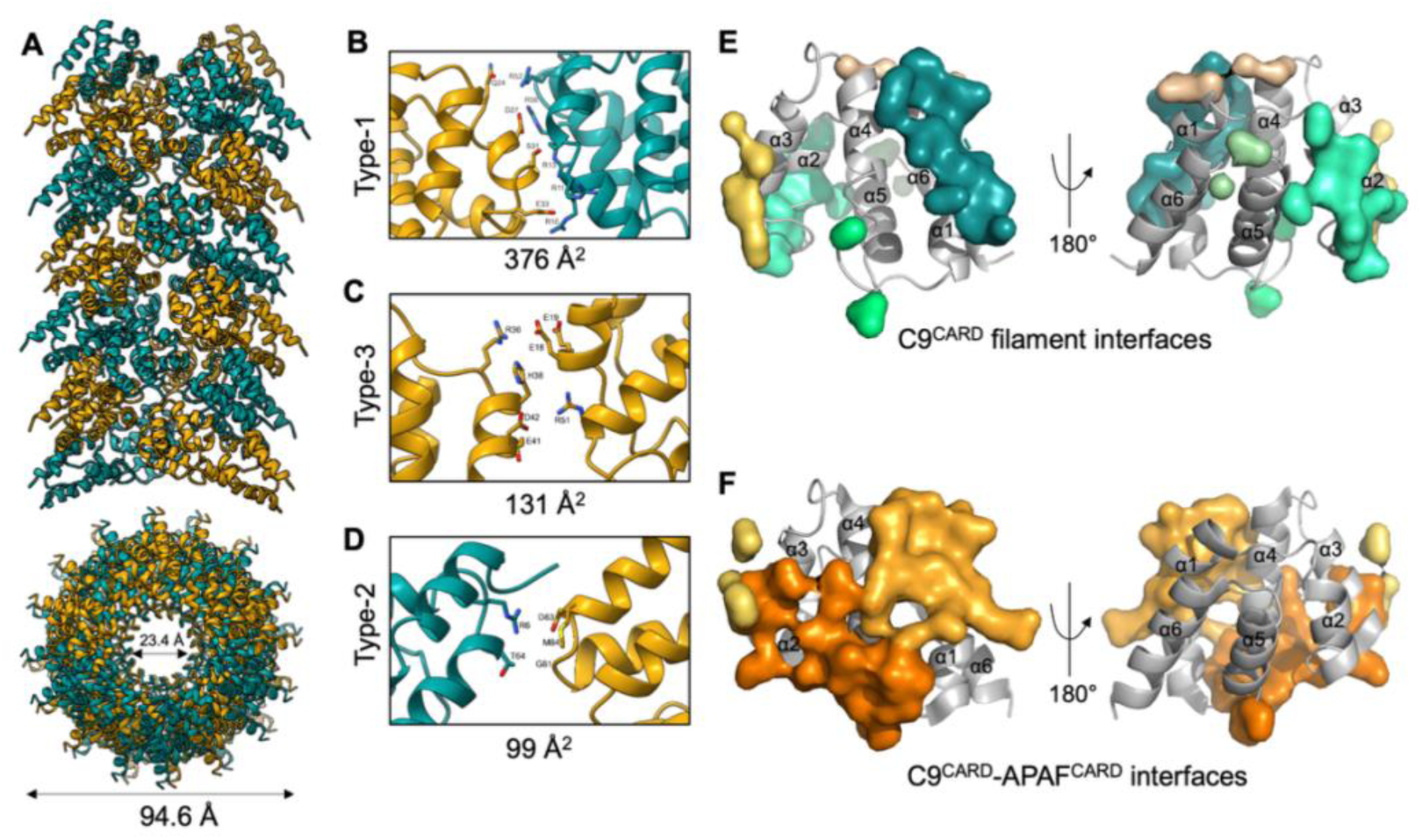
Cryo-EM structure of the C9^CARD^ filament. (**A**) Side view of the cryo-EM structure of the C9^CARD^ filament, including a rotated view down the filament that depicts the inner and outer diameters. Zoomed-in regions that correspond to the type-1 (**B**), type-3 (**C**), and type-2 (**D**) interfaces are shown along with the corresponding interface area in square Å. Key amino acids that make inter-protomer contacts are listed, including H38 in panel C. (**E**) CARD-CARD interfaces in the C9^CARD^ filament are shown on a single monomeric unit in differently colored surfaces. The helices are indicated. (**F**) The same analysis as in (E) except here for the C9^CARD^-APAF^CARD^ oligomeric CARD-CARD disk (PDB: 5wve).

Similar to other DDF filaments, the C9^CARD^ filament involves three major asymmetric interfaces with opposing interacting surfaces designated as *a* and *b* (**Figure 2B-D**, **Supplementary Figure 7**), with the type-1 and -2 interfaces mediating intra-strand interactions between the helical turns, whereas the type-3 interface forms inter-strand interactions in the helical strand direction (**Figure 2B-D**). The type-1 interface is the most extensive due to electrostatic complementarity, covering a total buried surface area of 376 Å^2^ (**Figure 2B**), and thus likely contributes the most to the binding energy required for filament formation. The type-1 interface is the most extensive due to the charge complementarity (376 Å^2^), followed by the type-3 and type-2 interfaces that contribute around 131 Å^2^ and 99 Å^2^ of buried surface area, respectively (**Figure 2B-D**). The type-1 interface involves interactions between residues that are located in the α2 and α3 helices (*e.g.*, Q24, D27, S31, Q33) of one subunit and residues in the α1 and α4 helices (*e.g.*, R56, R52, R13, R11, R10) of the adjacent subunit. This interface is stabilized by three hydrogen bonds and five salt bridges within 4 Å (**Figure 2C**). The type-3 interface consists of one hydrogen bond and three salt bridges between the residues of α2–α3 loop and α3 helix of one subunit and the α4 helix and α1–α2 loop of the adjacent subunit. Interactions are primarily formed between the arginine guanidinium groups and the side chain of H38, which is predominantly protonated at pH 5.5, with side chains of nearby glutamate and aspartate residues (**Figure 2D**). The type-2 interface only has two hydrogen bonds that connect residues in the α5–α6 loop and α6 helix of one chain to the α1 helix of the neighboring chain (**Figure 2D**).

We compared the molecular interactions within the C9^CARD^ filament to those formed with the Apaf-1 CARD (APAF^CARD^) in the CARD-CARD disk (PDB: 4rhw) (**Figure 2E, 2F**). Two APAF^CARD^ molecules interact with one C9^CARD^, with an overall stoichiometry in the crystal of 4:2. However, the interfaces in the APAF^CARD^:C9^CARD^ complex are not identical with those from our cryo-EM structure of the WT C9^CARD^ filament (**Figure 2F**). In complex with APAF^CARD^, the type-1 and -2 interfaces involve contacts between C9^CARD^ and APAF1^CARD^ whereas the type-3 interface derives solely from APAF^CARD^. Similar to the cryo-EM structure of the C9^CARD^ filament, the type-1 interface between residues of α1- α4 of C9^CARD^ (*e.g.*, R13, R52) and the α2-α3 of APAF^CARD^ (*e.g.*, D40, D27) shows multiple electrostatic interactions. Notably, the R52-D27 salt bridge is also observed in the cryo-EM structure of the WT C9^CARD^ filament. However, the type-1b interface of the filament involves residues in the α1 and α4 helices, which form the primary interaction surface for APAF^CARD^ (PDB: 3ygs), suggesting that the C9^CARD^ filament would be unable to associate with APAF^CARD^ or the apoptosome.

### Protonation of H38 promotes filament assembly

Having determined a high-resolution structure of the C9^CARD^ filament (**Figure 2A**), we next sought to better understand the mechanism of filament assembly. We observed that the C9^CARD^ more readily polymerizes into filaments at acidic pH. Moreover, the sequence of C9^CARD^ contains only a single His residue (H38) with an expected p*K*_a_ near physiological pH, while it is otherwise abundant in charged residues, which comprise nearly 33% of its sequence and are equally divided into positively charged (15 Arg, 1 Lys) and negatively charged residues (9 Asp, 7 Glu) (**Supplementary Figure 8**). Thus, we measured the solubility of the C9^CARD^ as the solution pH varied from 7 to 5.5 in 0.5-unit increments. The saturation concentration (*C*_sat_), or the maximum solubility of the protein given the solution conditions, reflects the concentration above which aggregates and filaments form (**Figure 3A**). While the C9^CARD^ remains highly soluble at neutral pH where the *C*_sat_ is 1.2 ± 0.1 mM, we observed a non-linear dependence of *C*_sat_ on pH, decreasing to 0.16 ± 0.01 mM at pH 5.5 (**Figure 3A**). As compared to the isolated CARD domain, both the C9^CARD+linker^ and full-length C9^C287A^ constructs were more soluble (**Supplementary Figure 9**), suggesting that the CARD-p20 linker and the protease domain modulate the propensity to form filaments (**Supplementary Figure 6**).

**Figure 3.**
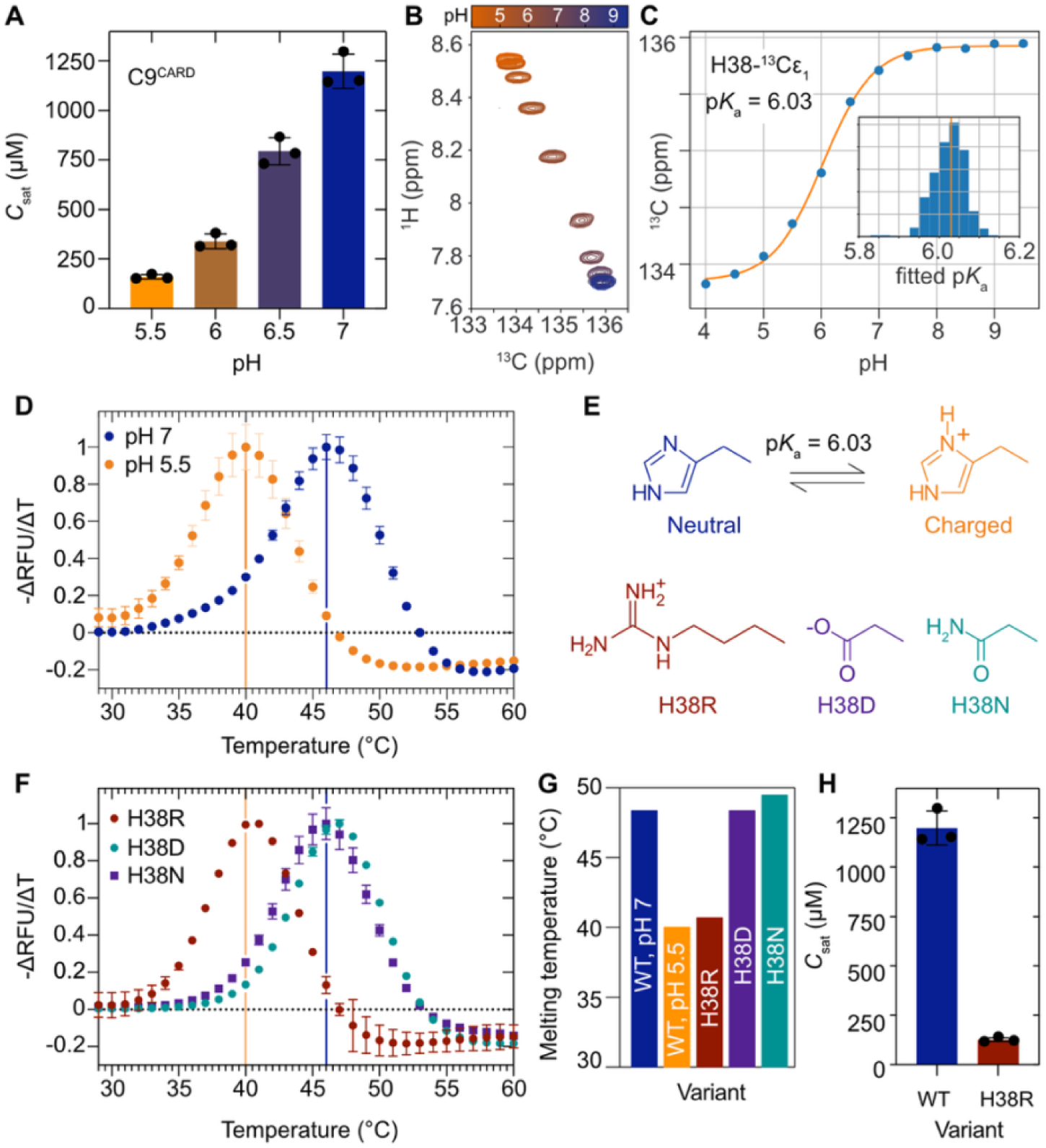
H38 protonation and the H38R mutation destabilize the CARD and enhance filament assembly. (**A**) *C*_sat_ measurements for C9^CARD^ as a function of the pH at 25 °C. (**B**) Overlaid 2D ^1^H-^13^C HSQC spectra of [U-^13^C,^15^N]-C9^CARD^ shown as a function of the pH, focusing on the side-chain ^13^Cε1 chemical shift of H38. (**C**) Variation in the H38 ^13^Cε chemical shift as a function of the pH, with the solid line corresponding to a fit to the Henderson-Hasselbalch equation. The inset shows the distribution of p*K*_a_ values obtained from a bootstrap error analysis, with the orange line positioned at the mean value of 6.03. (**D**) DSF-based thermal melt of C9^CARD^ in 25 mM HEPES, 2 mM DTT at pH 7 (blue) or pH 5.5 (orange). The normalized, negative derivative of the change in relative fluorescence units (RFU) with temperature is shown on the y-axis. (**E**) Equilibrium between the neutral and positively charged forms of H38, with the NMR-determined p*K*_a_ value indicated. Chemical structures of the side chains used to create variants at position 38: H38R (red), H38D (purple), H38N (green). (**F**) DSF-based thermal melt of C9^CARD^ variants H38R (red), H38D (green), and H38N (purple). The curves are compared to the WT C9^CARD^ at pH 7 (blue line) or pH 5.5 (orange line). (**G**) The melting temperatures for each variant. (**H**) *C*_sat_ determination for H38R at pH 7 and 25 °C, compared to the WT C9^CARD^ (same data from panel A).

Our observations suggest that the protonation of one or more titratable side chains between pH 7 to 5.5 enhances the propensity of the C9^CARD^ to self-assemble into filaments. Among its titratable side chains, the C9^CARD^ contains a single histidine residue (H38) with a theoretical p*K*_a_ of 6.5 that falls within the pH range described above. However, p*K*_a_ values can be dramatically shifted away from their theoretical values by local interactions, including electrostatics^86–88^, and as mentioned above the C9^CARD^ contains 33% charged residues. For this reason, we first used PROPKA^89^ to perform structure-based p*K*_a_ predictions on all available C9^CARD^ structures (PDB/chain: 4rhw/E, 3ygs/P, 5wvc/B, AlphaFoldDB/A, 5juy/O, 5wve/S), which returned a mean p*K*_a_ for H38 of 6.7 ± 0.4 (6.61, 6.20, 7.33, 6.15, 6.82, and 6.82, respectively) **(Supplementary Figure 10)**.

We then used NMR spectroscopy to monitor pH-induced changes to the ^13^C_ε1_ chemical shift of H38 (**Figure 3B**) and experimentally determine its p*K*_a_ (**Figure 3C**). Quantitative analysis revealed an apparent p*K*_a_ of 6.03 ± 0.04 (**Figure 3C**), which indicates that H38 is, indeed, predominantly protonated at pH 5.5 where the C9^CARD^ exhibits enhanced filament propensity, with an expected population of approximately 77 ± 1%. In addition, the C9^CARD^ is thermodynamically destabilized at pH 5.5, where it melts at 40 °C compared to 47 °C at pH 7 (**Figure 3D**), indicating that H38 protonation destabilizes the folded state.

### H38R mutation mimics the low-pH state and triggers filament assembly

Given that H38 is predominantly protonated at pH 5.5 where the C9^CARD^ is highly filamentous, we next evaluated how the stability and solubility are impacted by altering the electrostatic properties at this position (**Figure 3E**). To this end, we created three variants of the C9^CARD^ in which we altered the residue at position 38 to be positively charged (H38R), negatively charged (H38D), or polar but not titratable (H38N) (**Figure 3E**). We then measured the melting temperature and solubility of each variant at neutral pH (**Figure 3F**, **Figure 3G**). We observed that the H38R variant (T_m_ = 41 °C) was destabilized relative to the wild-type C9^CARD^ (47 °C) as well as the H38D (47 °C) and H38N (49 °C) variants (**Figure 3F**, **Figure 3G**). Indeed, the melting temperature of the H38R variant at pH 7 (41 °C) resembles that of the wild-type C9^CARD^ at pH 5.5 (40 °C) (**Figure 3F**, **Figure 3G**), which further indicates that a positive charge at residue 38 is destabilizing. By contrast, the H38D and H38N variants melt at the same or higher temperature as the wild-type C9^CARD^ (**Figure 3F**, **Figure 3G**).

We observed similar trends with protein solubility: at neutral pH, the H38R variant exhibited a *C*_sat_ of 130 ± 10 μM, which is nearly 10-fold lower than the WT C9^CARD^ at 1,200 ± 70 μM (**Figure 3H**). Negative-stain EM images confirm that the aggregates of H38R C9^CARD^ are filaments (**Supplementary Figure 11**), indicating that the H38R variant is highly filamentous, even at neutral pH. By contrast, H38N remained soluble up to protein concentrations that were more than 20-fold higher than the wild-type C9^CARD^, with no aggregates of H38N detected in this experiment. For H38R, the filament-enhancing effect can likely be attributed to the positive charge of the Arg head group and suggests that position 38 plays a key role in the filament assembly process. Using negative-stain EM, we observed filaments of H38R that resembled those formed by the wild-type C9^CARD^ (**Supplementary Figure 11**). Similarly, we observed filaments for the H38D variant that only formed at significantly higher protein concentrations as compared to the H38R variant (**Supplementary Figure 12**), suggesting that altering the residue at this site modulates but does not inhibit filament formation. Collectively, our results indicate that a positive charge at position 38 thermodynamically destabilizes the CARD, lowers its solubility, and promotes self-assembly into the filamentous state. These results support the notion that H38 is the primary pH sensor in the range of 7 to 5.5, where its protonation (or H38R mutation) contributes to polymerization.

### Cryo-EM structure of the H38R variant

The H38R filament bundles were long and straight (**Supplementary Figure 13**), and we determined a 3.5-Å structure of the H38R C9^CARD^ filament via cryo-EM (**Figure 4A**). Similar to the WT filament structure (**Figure 2A**), the tertiary structure of the H38R C9^CARD^ variant is preserved, with a backbone RMSD of 0.6 Å compared to the crystal structure of C9^CARD^ (PDB: 4rhw). The H38R filament has a twist angle of -67.2° and an axial rise of 9.1 Å per subunit, with an inner and outer diameter of 2.1 and 9.1 nm, respectively. These values differ slightly from the WT filament, leading to a subtle but significant altering of the packing and interfaces, with the result that all three types of interfaces in the H38R variant bury slightly more surface area than the in the WT filament.

**Figure 4.**
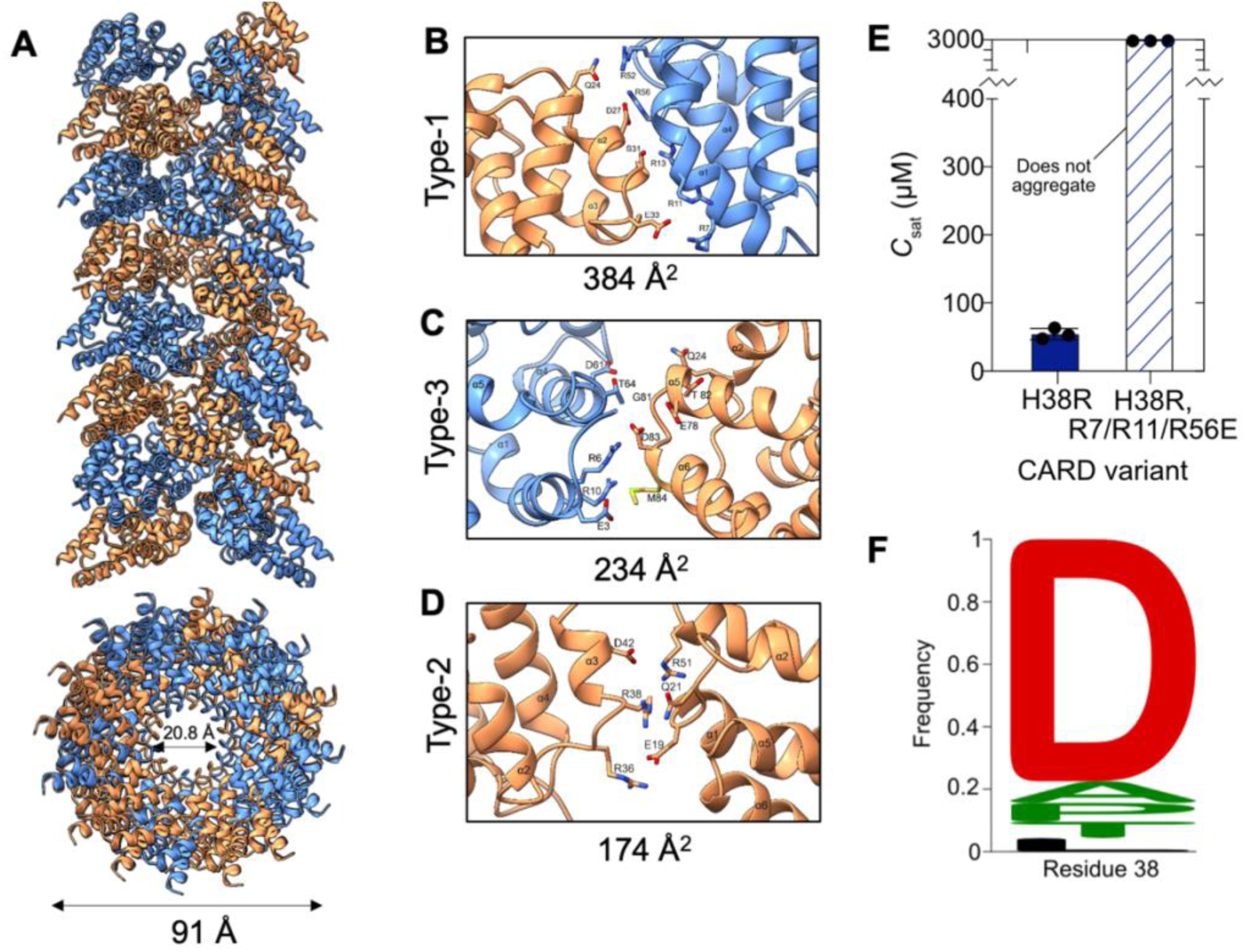
Cryo-EM structure of the H38R C9^CARD^ filament. (**A**) Side view of the cryo-EM structure of the H38R C9^CARD^ filament, including a rotated view down the filament that depicts the inner and outer diameters. Zoomed-in regions that correspond to the H38R type-1 (**C**), type-3 (**D**), and type-2 (**E**) interfaces are shown along with the corresponding interface area in square Å. (**E**) *C*_sat_ measurements for the H38R and H38R triple mutant (R7/R11/R56E) at pH 5.5 and 25 °C. These mutations were designed to disrupt the type-1 interface of the filament. (**F**) Sequence logo of residue preferences at position 38 (human numbering) in C9 across 265 different orthologs. Residues are colored by hydrophobicity and are shown with 10 or more hits: D = 179, A = 16, P = 14, T = 12, and L = 10.

For the H38R filament, the type-1 interface mediates interaction between residues in the α2 and α3 helices (e.g., E33, S31, D27, Q24) of one subunit with residues in the α1 and α4 helices (e.g., R52, R56, R13, R11, R7) of the neighboring subunit (**Figure 4B**). Within a distance of 4 Å, there are a total of six hydrogen bonds and four salt bridges contributing to this interface. The type-3 interface (**Figure 4C**) buries 234 Å^2^ of surface area, contains R38 (via H38R) as well as salt bridges between the guanidinium group of arginine and side chains of glutamate and aspartate. In total, there are four salt bridges and three hydrogen bonds that contribute to this interface. The type-3 interface mediates interaction between residues in the α2-α3 loop and the α3 helix of one subunit with the residues in the α4 helix and α1-α2 loop of the neighboring subunit. The type-2 interface, with a total buried surface area of 174 Å^2^ (**Figure 4D**), is the least extensive among all three interfaces. It constitutes only two hydrogen bonds between residues in the α5-α6 loop and α6 helix of one chain to the α1 helix of the neighboring chain.

To test the cryo-EM model of the H38R C9^CARD^ filament, we generated a triple-mutant in which the mutations R7E, R11E, R56E were introduced to the H38R variant (**Figure 4E**). These charge-reversing mutations were designed to disrupt interactions across the type-1 interface of the C9^CARD^ filament. We purified the H38R triple-mutant and measured its solubility under filament-forming conditions at pH 5.5 where the *C*_sat_ of H38R is approximately 50 μM. However, we were not able to induce aggregates of the H38R triple-mutant, suggesting that it is unable to form filaments even at high protein concentrations above 3 mM (**Figure 4E**). Thus, the structure-based mutations that we introduced into the H38R variant raised its *C*_sat_ by more than 60-fold, from *ca.* 50 μM to greater than 3 mM.

To better understand the sequence properties at the H38 position in C9, we next investigated the natural sequence variation across more than 250 different C9 orthologs. The most common residue type at this position is the negatively charged D (*n* = 179), followed by A (16), P (14), T (12), L (10), H (7), N (7), G (6), and S (6) (**Figure 4F**). We did not find a positively charged residue at this position in any of the examined organisms. Interestingly, the H38R variant in C9 is present as a low-frequency single-nucleotide polymorphism in the human population, as reported by the gnomAD database^90^. The pathological significance of this variant, if any, is not presently known. Our sequence analyses indicate that a negatively charged residue at residue 38 (human numbering) is preferred in the C9^CARD^.

### Protonation of H38 alters the populations of side-chain rotamers

Our data indicate that C9^CARD^ polymerizes into filaments through a process that is favored by H38 protonation or the H38R mutation. Moreover, H38 protonation event or the H38R mutation also destabilize the CARD fold, lowering the melting temperature from 47 to 40 °C or 41 °C, respectively. From a structural perspective, however, it is unclear why the H38R mutation or H38 protonation destabilize the domain. Inspection of the C9^CARD^ structure (PDB: 4rhw) shows that H38 is a solvent-exposed residue that is next to (< 3.5 Å) two negatively charged residues (E41, D42). Indeed, the structure-based predictors of protein stability FoldX^91^ and DDmut^92,93^ do not predict any significant changes to the stability of the C9^CARD^ following the H38R mutation (ΔΔG = -0.19 kcal mol^-1^ vs. RMSE of 1.37 kcal/mol on cross-validation set).

We hypothesized that H38 protonation or the H38R mutation may be destabilizing due to electrostatic incompatibility with the helix dipole, which produces a partial positive charge at helical N-termini^94^. H38 is the second residue of α-helix 3 in the C9^CARD^, referred to as N2 using the Richardson and Richardson terminology^95^. In an α-helix, a partial positive charge forms at the N-terminus due to the parallel orientation of backbone amide N-H bond vectors^96–98^. By contrast, a partial negative charge forms at the C-terminus because of the alignment of backbone carbonyl C-O bond vectors. To better understand the destabilizing effects of H38 protonation and the H38R mutation, we performed all-atom molecular dynamics (MD) simulations of the C9^CARD^ as a function of the H38 protonation state, or with the mutation H38R (**Figure 5, Supplementary Figure 14**).

**Figure 5.**
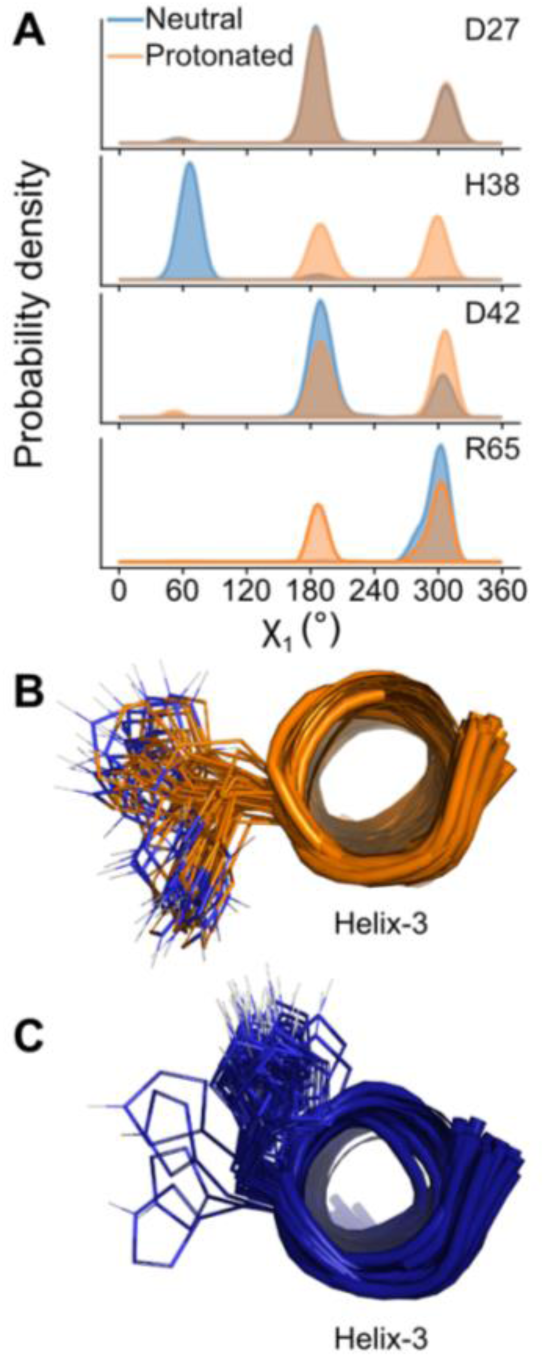
Protonation of H38 alters its side-chain rotamer populations. (**A**) χ_1_ rotamer populations (*gauche*+, *trans*, *gauche-*) of the selected side chains derived from the 5-μs MD simulations of the H38-neutral (blue) or H38-protonated (orange) forms of the C9^CARD^. (**B**, **C**) Snapshots of helix-3 from the MD trajectories of the C9^CARD^ in the H38-protonated (**B**) or H38-neutral (**C**) form with the side-chain of H38 shown in stick format. H38 protonation orients the imidazolium group toward solvent whereas the neutral form is oriented toward the N-terminal helix dipole.

In the structure of C9^CARD^ (PDB: 4rhw, chain E), which was crystallized at pH 5.5, the side chain of H38 is surrounded by nearby negatively charged residues in helix-3, forming interactions with the side chains of residues that are i+3 (E41, 3.0 Å) and i+4 (D42, 3.4 Å). The acidic pH of the crystallization buffer^72^ suggests that H38 was likely protonated. Our MD simulations of the C9^CARD^ with H38 in either the neutral or protonated form show that the global fold of the domain is preserved over the course of the trajectory (**Supplementary Figure 14**). H38 protonation does not impact the per-residue root-mean-square fluctuation (RMSF) values for most residues (± 0.1 Å compared to neutral-H38) (**Supplementary Figure 14**) and a few sites with larger deviations. For H38 itself, there is no difference in backbone RMSF for the neutral or protonated state. However, among side-chain heavy-atom RMSF values, H38 shows the second largest difference in RMSF values (**Supplementary Figure 14**) (neutral – protonated = -0.5 Å), pointing to increased structural dynamics in the protonated form.

Consistent with the above RMSD and RMSF analyses, we found that H38 protonation does not cause any major changes to the backbone ɸ and ψ dihedral angle distributions that were observed in the H38-neutral state. Instead, we observed large redistributions to some χ_1_ side-chain rotamer populations, including H38 itself (**Figure 5A, 5B, 5C, Supplementary Figure 15**). In its neutral form, H38 almost exclusively populates the *gauche+* χ_1_ rotamer (60°; *ca.* 98% population), which is the rarest His rotamer and populated to less than 15% among His residues in the Dunbrack rotamer library^99^. In this rotamer, the side chain of H38 is oriented toward helix-3 of the C9^CARD^. Interestingly, a statistical analysis of helical residues in the PDB identified that His in the N2 position is nearly 2-fold enriched in the *gauche*+ χ_1_ rotamer compared to the background distribution of all other His residues; by contrast, His residues in helical interiors (N4 to C4 positions) almost never adopt the *gauche*+ rotamer^100^.

Upon protonation, H38 undergoes complete conversion to the more common *trans* (180°; 48%) and *gauche-* (300°; 52%) χ_1_ rotamers (**Figure 5A**). Both χ_1_ rotamers reposition the imidazolium of H38 away from helix-3 and toward solvent (**Figure 5B**). These observations indicate that protonation of H38 induces a transition from the rare *gauche+* rotamer, with the side chain oriented towards helix-3 (**Figure 5C**), to the more common *trans* and *gauche-* rotamers in which the side chain now points toward solvent and away from the helix dipole (**Figure 5B**).

For the H38R variant, our MD simulations show that this mutation does not alter the global structural and dynamical properties of the C9^CARD^. However, at a more local level, we observed that the R38 side chain populates rare regions of Arg-rotamer space (**Supplementary Figure 16**). For example, in [χ_1_, χ_2_] space, the R38 side chain predominantly samples [*gauche-, gauche-*] followed by [*trans*, *gauche+*] rotamers. Among all other 15 Arg residues in C9^CARD^, the major [χ_1_, χ_2_] conformers are [*gauche-*, *trans*] and [*trans, trans*]. Similarly, in [χ_3_, χ_4_] space, the majority of R38 conformers are near (*gauche+, gauche+*) as compared to [*trans, trans*] for all other 15 Arg residues (**Supplementary Figure 16**). Thus, the most-abundant R38 side-chain rotamers for [χ_1_, χ_2_, χ_3_, χ_4_] are [300°, 300°, 60°, 60°] and [180°, 60°, 180°, 180°] compared to [300°, 180°, 180°, 180°] among the other Arg residues. These unique χ_1-4_ combinations respectively place the R38 headgroup in a favorable position to form a salt bridge with the side chain of E41, located *i*+3 to R38, or to maximize the distance between the R38 headgroup and the helix dipole (**Supplementary Figure 16**).

Therefore, our MD simulations of the C9^CARD^ in H38-neutral or H38-protonated forms, as well as with the H38R mutation, show that a positive charge at this residue leads to sampling of rotamers that increase the distance between the helix dipole and the side chain. In the case of H38R, a salt bridge to the i+3 helical residue (E41) transiently forms, although this interaction is not stably formed and the R38 side chain repeatedly interconverts between rotameric states that place the Arg head group away from the helix dipole.

### Survey of helical protein filaments reveals frequent histidine placement at interfaces

Our results above indicate that the pH-regulated filament formation of C9^CARD^ manifests due to a fine-tuned balance between competing, pH-dependent effects: intramolecular destabilization (unfolding) and intermolecular contacts (self-assembly). On the one hand, protonation of H38 thermodynamically destabilizes the domain due to an electrostatic clash between the cationic imidazolium and the partial positive charge of the N-terminal helix-3 dipole, lowering the melting temperature by 7 °C. On the other hand, H38 protonation induces a χ_1_ rotameric transition that orients its side-chain toward solvent, where it can mediate inter-molecular contacts that contribute to self-assembly.

How many other protein filaments may be regulated by pH in a similar manner? We searched specifically for helical protein filaments that contain His at the filament interface. To this end, we curated a database of 173 helical protein filaments from the PDB (**Methods**), excluding amyloid fibrils. We redundancy filtered this set, yielding a total of 77 non-redundant filaments. We then extracted all interfacial residues in each structure and identified 30 filaments that contain His at one or more interfaces, including NINJ1 (8qcr), RAD51 (8bsc), *B. subtilis* flagella (5wjt), actin (6tu4, 7q8c, 8ccn, 6iug), NLRP6 PYD (6ncv), tankyrase-2 (8aly), and Acetyl-coenzyme A synthetase (8rwj), among others (**Supplementary Table 1**). However, the mere presence of a His residue at a filament interface does not *per se* confer pH-regulated self-assembly, as evidenced by the lack of pH-dependent assembly in human CTP synthase^32^. Rather, the placement of histidine residues near nearby electrostatic groups, *e.g.* helix dipoles or charged amino acids, is likely required.

### Proteome-wide identification of pH-sensitive histidine residues near helix dipoles

This prompted us to ask how many other protein domains may contain a pH-regulated histidine switch near an N-terminal helix dipole. Mapping such sites could inform on other, endogenous pH sensors that regulate molecular recognition, self-assembly, or conformational transitions more broadly. This has been observed in other oligomers, for example, in tetradecameric ClpP protease whereby the protonation of a single His residue located *i-1* to an N-cap shifts a conformational equilibrium from the active “extended” to the inactive “compressed” conformation^101^. Therefore, we leveraged the recently developed repository of AlphaFold2-predicted domains for the human proteome (AF2 ECOD)^102^ (**Figure 6A**) to search for those that contain His near an N-terminal helix dipole. We assigned the secondary structures with DSSP to obtain more than 140,000 helices of suitable length (**Methods**). This represents our AF2-ECOD helix residue atlas (**Figure 6A**), which comprises a significantly larger dataset than earlier, pioneering studies that were limited by the size of the Protein Data Bank at that time^95,100,103^. We mined the AF2-ECOD helix residue atlas to identify protein domains that contain a His residue near an N-terminal helix dipole, where the His residue may act as an endogenous pH sensor, either by decreasing protein stability or by functioning a molecular “switch”.

**Figure 6.**
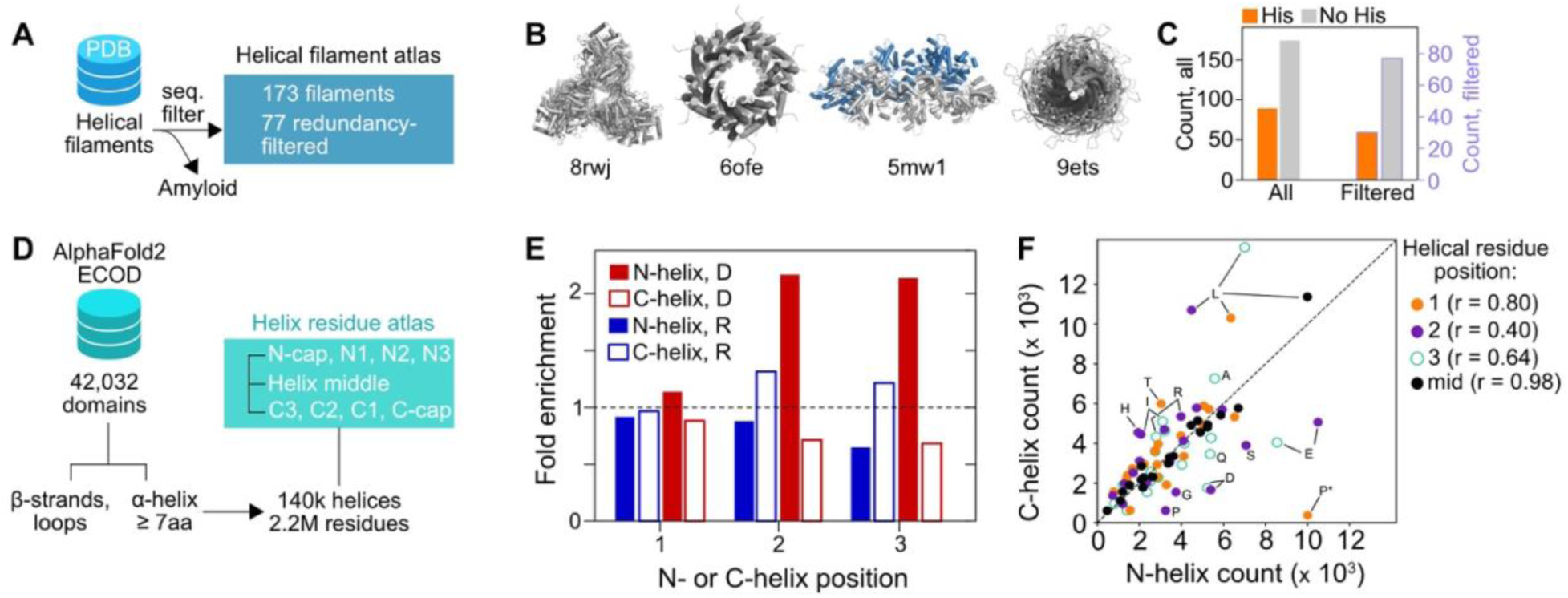
Atlases of helical filaments and helical residue preferences. (**A**) Helical filaments were extracted from the PDB, ignoring amyloid fibril structures, and redundancy-filtered for similar sequences. (**B**) Example structures of helical filaments with the corresponding PDB codes listed. (**C**) For the 173 (77) helical filaments in the atlas (redundancy filtered atlas), a total of 89 (30) contain one or more His residue at a filament interface. (**D**) Flowchart of the AF2 ECOD^102^ helical residue atlas. (**E**) Fold enrichment of R and D residue types at the positions N1/C1, N2/C2, or N3/C3 in the AF2 ECOD helix residue atlas. N-terminal (C-terminal) helix positions are shown in filled (empty) bars. See Methods for the definition of fold enrichment. (**F**) Correlation of N-terminal helix residue positions (x-axis) with C-terminal helix residue positions (y-axis) for the N1/C1 (orange), N2/C2 (purple), and N3/C3 (green, empty) positions. Residues in the middle of helices are shown in black. The Pearson correlation coefficient is indicated, and residue types far from y=x are labeled.

Of the *n* = 42,032 domains in AF2-ECOD, there are 31,712 domains that contain an α-helix of seven or more residues in length. We identified a total of *n* = 7,883 (24.9%) domains that contain at least one His residue in at least one of the N-cap, N1, or N2 positions of a helix. When analyzed by His-residue count, we found that 76% of the 7,883 domains contained only a single His residue at the N-cap, N1, or N2 position (**Supplementary Table 2**). Overall, there are 307 different domain families that harbor His near an N-terminal helix dipole (**Supplementary Table 2**), indicating a diverse landscape of potentially pH-regulatable protein domains. Examples domain types include zinc fingers, AAA+ ATPase lids, TIM barrels, RING / U-box, EF-hand, FF, POZ, PAH2, SH3, and PDZ, among others (**Supplementary Table 2**). This represents only a minor fraction of the 1,661 human domain families (18.5%).

Of the 1,857 domains (24%) that contained more than one His residue near an N-terminal helix dipole, the top 10 hits, which contain more than 10 His residues each, are repeat domains that contain either Ankyrin or ARM repeats. For example, the top hit (ANKRD17, UniProt O75179) is an Ankyrin repeat protein that contains 17 His residues that are all at an N-cap position. Such repeat domains may be particularly sensitive to pH changes, given that these domains contain multiple His residues positioned near helical N-terminal dipoles.

To examine the residue frequency in helices more generally, we leveraged the recently developed repository of AlphaFold2-predicted domains for the human proteome (AF2 ECOD)^102^ (**Figure 6A**). For the *n* = 42,032 ECOD-classified domains in the human proteome, we assigned the secondary structures with DSSP to obtain more than 140,000 helices of suitable length (**Methods**). This represents a significantly larger dataset than early, pioneering studies that were limited by the size of the Protein Data Bank at that time^95,100,103^. We then compared residue types by position in helices (**Figure 6A**). Specifically, we compared residue types at helical N-termini (N1, N2, N3) versus those in the corresponding positions at helical C-termini (C1, C2, C3). The corresponding analysis for N- and C-cap residues is included in the Supplementary Information (**Supplementary Figure 17**). Normalized to all helical residues in the human proteome, we observe a strong enrichment for D in the N2 and N3 positions, but not for the N1 position (**Figure 6B**). By contrast, we observe a strong depletion of R in N3 position, with a mild depletion at N2 and little difference at N1. These results indicate that, indeed, D is preferred near the N-termini of helices while R is not (**Figure 6B**, filled bars). At helical C-termini, however, the situation is reversed: D is strongly depleted, especially at C2 and C3 positions, while R is enriched at C2 and C3 (**Figure 6B**, empty bars). When comparing the residue-type counts between N- and C-helix positions across all residues (**Figure 6C**), we find that the highest Pearson correlation coefficient is obtained for residues that approach the helix middle (r = 0.98, black), whereas the N2/C2 and N3/C3 comparisons yield significantly more deviant relationships (r = 0.40, purple and r = 0.64, green) (**Figure 6C**). The N1/C1 comparison, however, is more correlated with r = 0.80 (**Figure 6C**, orange), suggesting more similar residue-type usage, which is also reflected in the lower levels of deviation in the R vs. D comparisons for the N1 and C1 positions (**Figure 6B**). Interestingly, P is the most abundant N1 residue type (*n* = 10,035 or 13.7%) followed by E (6,525 or 8.9%). In the case of P, this residue does not contribute a free N-H group and therefore does not constructively add to the helix dipole, whereas E provides a negatively charged side chain that can interact favorably with the helix dipole.

### Widespread destabilization by positively charged residues at the N-termini of α-helices

The helix dipole effect on protein stability has been explored in a handful of model peptides and proteins^96–98,–107^; however, the impact in other proteins at a broader scale has not been investigated. Thus, to examine the impact of mutations near helical termini in hundreds of different protein domains, we turned to a recently developed deep mutational scanning (DMS) resource^108^ known as the Human Domainome 1. With over 600,000 mutations in more than 500 distinct protein domains, the Human Domainome 1 offers a rich coverage of the sequence-fitness landscape^108^. Notably, there are two CARDs within the Human Domainome 1 (**Figure 7A**, **Figure 7B**), enabling us to examine the effects of mutations near helix dipoles in a similar fold as the C9^CARD^. There are a total of 3,243 mutations and fitness measurements from the CARDs of Receptor-interacting serine/threonine-protein kinase 2 (RIPK2^CARD^, UniProt O43353) and Apoptosis-associated speck-like protein containing a CARD (ASC^CARD^, UniProt Q9ULZ3). In RIPK2^CARD^, the N1 residues of helix-2 and helix-5 are negatively charged, and charge-reversing mutations at these sites significantly decrease fitness (mean value = -0.75 ± 0.12 [E453K = -0.54, E453R = -0.79; E499K = - 0.80, E499R = -0.86]), with a magnitude that is comparable to the effect of a nonsense mutation (E453* = -1.17, E499* = -1.06) (**Figure 7C**, black). In ASC^CARD^, helix-3 contains only a single negatively charged N1 residue (D143). Charge-swapping mutations at this site destabilize the domain (D143K = -0.42, D143R = - 0.25), although to a lesser extent than observed in RIPK2^CARD^ (D143* = -0.91) (**Figure 7C**). Similarly, in both CARDs we observed a decrease in fitness upon charge-reversing mutations at N2 and N3 sites or introduction of positive charges at otherwise polar N-cap residues (**Figure 7C**).

**Figure 7.**
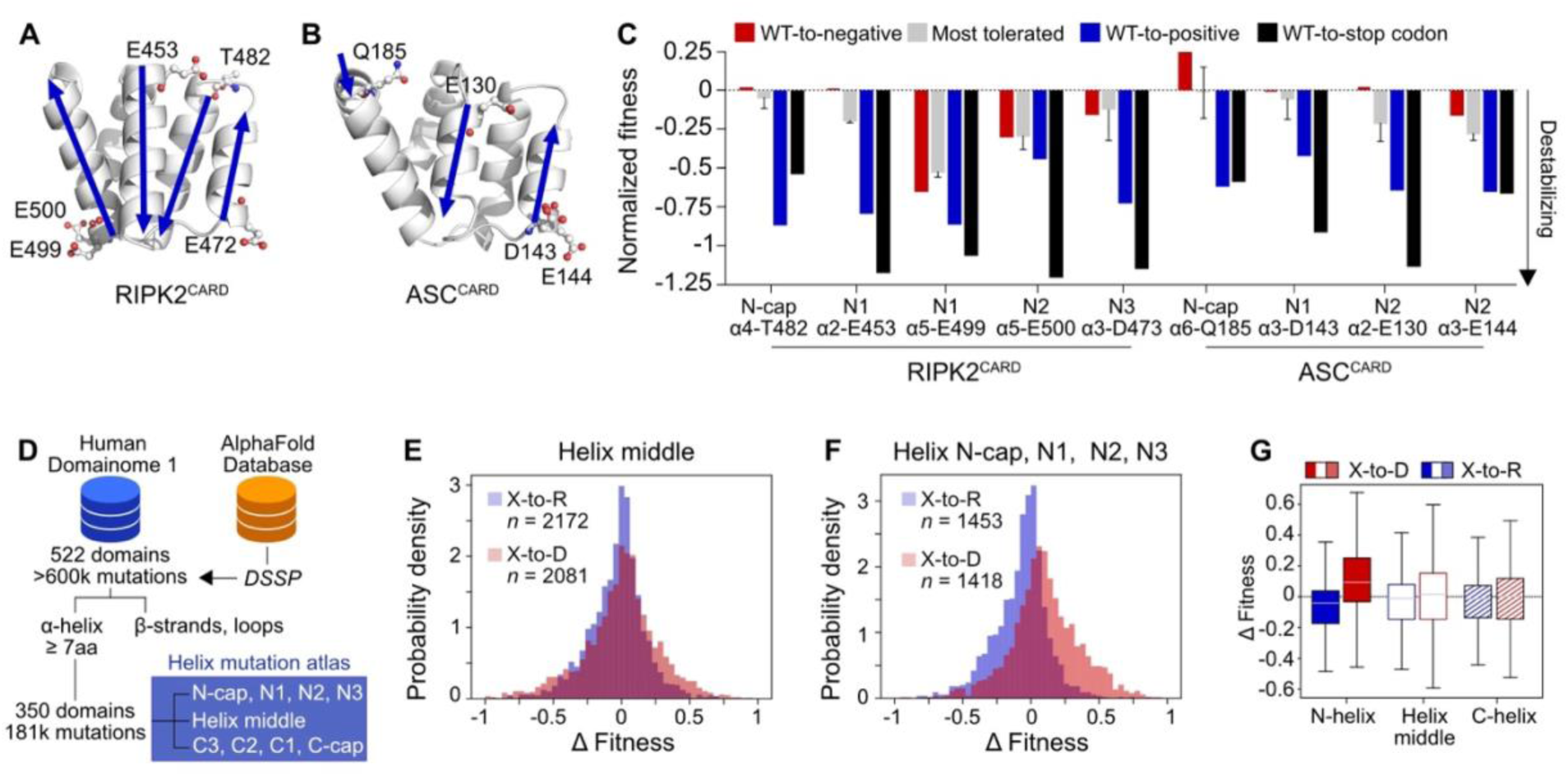
Destabilizing effects of charge swaps at N-terminal helix dipoles. (**A**) AlphaFold2-predicted structures of the RIPK2^CARD^ and (**B**) ASC^CARD^ as well as the negatively charged or polar residue types near the N-termini of the indicated helices. The helix directions (N-to-C) are shown with blue arrows. (**C**) The normalized fitness for the residues shown in panel D extracted from the Human Domainome 1 dataset^108^. Mutations to a negatively charged residue (i.e. D/E-to-E/D, or polar-to-D/E) are red. Mutations to a positively charged residue (i.e. D/E-to-R/K or polar-to-R/K) are shown in blue. The mean ± standard deviation of the three most tolerated mutations, excluding mutations to charged residues, are shown in grey. Nonsense mutations are shown in black. (**D**) Filtering of the Human Domainome 1 dataset to create the helix mutation atlas. (**E**) Distribution of the change in fitness (Δ Fitness, see Methods) upon mutation of a helix middle residue from X-to-R (blue) or X-to-D (red), where X is defined as a polar residue. (**F**) The same as (E) but for N-helix residues, i.e. N-cap, N1, N2, and N3 residues. (**G**) Box plots of the data in (E) and (F), and for C-helix residues. The median values are shown in the grey line and the boxes include data points between the first and third quartiles, with the whiskers extending to 1.5-fold the interquartile range.

Beyond helix dipole effects, the intrinsic “mutability”, or sensitivity to mutation at these sites, reflects the local structural context. For example, in RIPK2^CARD^ the conservative mutation E453D is effectively neutral (+0.01), with other mutations at this site that are only mildly destabilizing (E453N = -0.21, E453M = -0.19, E453T = -0.20) (**Figure 7C**). By contrast, the conservative mutation E499D destabilizes RIPK2^CARD^ (-0.65) and all other mutations approach the effect of a nonsense mutation, with the most tolerated mutations at this site (E499P/G/A, -0.49/-0.54/-0.55, *grey*) approaching a nonsense mutation (**Figure 7C**, black). In ASC^CARD^, D143 is more tolerant to mutations, including D143E (-0.01), D143A (-0.16), D143H (+0.09), D143L (-0.1). Local structural context thus plays an important role, as evidence by negative fitness impacts upon the removal of some positively charged N-cap residues. For example, in the RIPK2^CARD^, substituting two positively charged N-cap residues (K471 in helix-3; R483 in helix-4) with negatively charged residues decreased overall fitness (R483D = -0.22; K471D = -0.45). Thus, the introduction of positively charged residues at N-cap, N1, N2, or N3 positions in the RIPK2^CARD^ and ASC^CARD^ decreased fitness when compared to negative-to-negative mutations (i.e. E-to-D, **Figure 7C**, red) or the three most tolerated mutations at each position (**Figure 7C**, grey).

Next, we wondered if introducing positive charge near the N-terminus of helices generally decreases protein stability in other folds. We then expanded our analysis to include all α-helices within the Human Domainome 1^108^ (**Figure 7D**). After filtering (see Methods), we extracted 783 α-helices from 350 unique protein domains, totaling 9,521 residues for which 180,889 mutations are available (**Figure 7D**). This represents approximately 30% of the entire Human Domainome 1 dataset (*n* = 602,882 total mutations). The mean helix length in our filtered dataset is 12 ± 4 residues, as helices with fewer than two turns were removed. With our filtered helical Human Domainome 1 dataset, we then segmented each helix into the N-terminal or C-terminal residues (i.e., N-cap, N1, N2, N3 vs. C-cap, C1, C2, C3), or the helix middle. We extracted all mutations X-to-D and X-to-R, where X was defined as a polar or charged residue, and computed a helix-normalized fitness defect (**Figure 7E**, **Figure 7F**, Methods). The distributions of fitness defects show that X-to-R mutations at helical N-termini (*n* = 1,453) are significantly more destabilizing than X-to-D mutations (*n* = 1,418) (**Figure 7G**). By comparison, we did not see significant deviations in fitness defects for such mutations in helical C-termini (**Figure 7G**) or the middle of helices (**Figure 7G**), with only a marginal increase in the upper tail of the distribution for X-to-D mutations. This is consistent with previous reports of helical peptides tolerating more substitutions at their C-termini^109^.

Thus, introduction of R vs. D residues at the N-termini of helices is significantly more destabilizing than in the middle of helices or at the C-termini. Nonetheless, we found thousands of instances in which a protein domain contains R near a helical N-terminus (N-cap, or N1-3). We speculate that this may be allowed for one of two reasons: (1) the intrinsic stability of such domains may be higher, enabling toleration to a locally unfavorable interaction with the helix dipole or (2) the local structural context may position negatively charged residues in direct vicinity, e.g. at i+3 or i+4 in the helix to enable intra-helix salt bridges. The latter type of interaction has been recently described in highly thermostable, *de novo* designed protein domains, including an R at helix N-terminus that is involved in an i+4 salt bridge^110^.

## Discussion

Self-assembling helical protein filaments are common throughout biology and enable localized, rapid biochemical studies^54–58^ amplification of biochemical signals ^14,15 18–20 21,22^. More recently, engineered or designed helical filaments have been successfully used in various biotechnological applications ^24,25 26,27^. However, engineering proteins to form helical filaments in response to environmental cues remains challenging ^24,25,28 29^. Here, we found that a natural protein domain, the human C9^CARD^, polymerizes into helical filaments in a pH-responsive manner. The C9^CARD^ was not previously known to self-assemble, despite many structural and,^58–72,72,73,73–77^.

We dissected the solution conditions that drive C9^CARD^ self-assembly, finding enhanced filament assembly under low-salt or acidic conditions, which suggests a key role for inter-molecular electrostatic interactions. Charge-altering mutations of the lone histidine residue to introduce a positive (H38R) or negative (H38D) charge, or to remove the charge at this site altogether (H38N), dramatically modulated the self-assembly of the C9^CARD^, indicating that interactions involving H38 directly contribute to filament formation. We used cryo-electron microscopy to determine 3.3- and 3.5-Å structures of the WT C9^CARD^ and H38R filaments. Like other DDFs filament structures, C9^CARD^ shows extensive electrostatic interactions between the CARD subunits within the filament at the type-1 interface, while H38 is involved in the type-3 interface, and the H38R increases the size of this interface. The increased area of the type-3 interface in the H38R variant explains why this mutant is highly filamentous. Rationally designed mutations, based on our cryo-EM structure, were sufficient to disrupt filament formation in the otherwise highly filamentous H38R variant, increasing the *C*_sat_ by nearly two orders of magnitude.

Using NMR spectroscopy, we confirmed that the side chain of H38 is predominantly protonated at pH 5.5 where the C9^CARD^ is highly filamentous. Our NMR-determined p*K*_a_ value of 6.03 ± 0.04 for H38 in the N2 position agrees with studies in model peptides where a His residue in a nearby N-cap position yields a reduced p*K*_a_ of 6.1, due to the helix dipole, as compared to 7.4 in a C-cap position^93,97,111^. Furthermore, in agreement with our thermal melting data, the protonation of a His residue in an N-cap position was found to significantly destabilize helical peptides^112^, and protonated His was found to be among the least favorable N2 residue types in a study employing small helical peptides^113^. We further employed an integrative biophysical and structural approach to understand the destabilizing effect of H38 protonation or the H38R mutation, identifying an electrostatic clash with the partial positive charge from the helix dipole that leads to a rotameric switch to reposition the side-chains toward solvent. The destabilizing effect of protonated H38 or H38R is presumably neutralized upon filament assembly, in which H/R38 is positioned near two negatively charged residues, E18 and E19, as well as a C-terminal helix dipole that bears a partial negative charge. Therefore, His protonation regulates a fine balance between intramolecular destabilization (unfolding) and intermolecular contacts (self-assembly). This suggests that the intrinsic stability of a pH-regulated domain must be sufficiently high to “tolerate” the electrostatic destabilization.

The destabilizing effect of charged residues positioned near helix dipoles has been described in several monomeric proteins and short peptides. However, this effect has not been examined on a larger scale. We leveraged AlphaFold2-based structure prediction and the recently established Human Domainome 1 DMS resource, encompassing more than 150,000 protein stability measurements across more than 350 helix-containing protein domains, including two CARDs. Our results demonstrate that introducing positive charge near the N-terminal helix dipole significantly decreases protein stability. However, this effect vanishes when positively charged residues are introduced in the middle of helices or at the C-terminal helix dipole. With AlphaFold2-based protein structure prediction on a proteome scale, we further quantified the residue usage frequency in helices across all human protein domains. We observed a strong increase (decrease) in R (D) content near N-terminal helix dipoles, supporting our DMS analysis.

In summary, we identified a helical protein filament that is formed by the C9^CARD^ and investigated the biophysical properties that drive filament formation. We identified a histidine residue (H38) whose protonation or Arg-mutation stimulates filament assembly, thus functioning as a pH-sensitive switch that regulates CARD polymerization. Indeed, removing the charge at this position via the H38N mutation generated a highly soluble protein whose *C*_sat_ could not be determined. Cryo-EM structures of the wild-type and H38R C9^CARD^ filaments reveal a similar overall architecture to other CARD filaments and expand the catalog of known DDF filaments. Our identification of a natural, pH-responsive protein filament may be relevant for the *de-novo* design of protein filaments: Shen and colleagues have recently described an approach to design pH-responsive protein filaments that leverage His protonation^25^. Their designs require extensive networks of buried histidine residues, whereas the C9^CARD^ contains only one. Taken together, our findings suggest that C9^CARD^ polymerization is finely tuned by the pH through a single histidine residue that modulates charge at a structurally sensitive site. Together with our broad-scale AlphaFold2 and DMS analyses, our results point to an electrostatic design principle in which charge placement near N-terminal helix dipoles could be leveraged to tune side-chain rotamers and self-assembly for environmentally responsive filaments.

## Methods

### Protein expression and purification

DNA encoding for human caspase-9 (UniProt ID: P55211) was synthesized and codon optimized for *E. coli* expression by GenScript^63^. The gene was then sub-cloned into a pET-29b(+) expression vector that contained an N-terminal hexahistidine-tag followed by small ubiquitin-like modifier (His-SUMO) solubility tag. Two fragments of caspase-9 were derived from the full-length plasmid via mutagenesis: the CARD domain (residues 1-99) and the CARD+linker (residues 1-138)^63^. All mutagenesis, including point mutants of H38 (vide infra) and elsewhere in the CARD, was performed by GenScript. All of the caspase-9 expression constructs contain a Ser-Gly linker that separates His-SUMO from Met1 of the CARD domain. This was found to be necessary to enable complete cleavage of the His-SUMO tag by the Ulp1R3 SUMO protease^63^, likely for steric reasons due to the burial of Met1 in the structure of the CARD.

For recombinant protein expression and purification, we followed previously established protocols for C9^CARD^+linker and C9 (C287A) ^63^. All variants were expressed and purified similarly. For the production of ^15^N-labeled or ^13^C,^15^N-labeled protein, M9 minimal medium was used with 1 g/L of ^15^NH4Cl and 6 g/L of D-glucose or 1 g/L of NH_4_Cl and 2 g/L of uniformly ^13^C-labeled D-glucose, respectively. Protein purification proceeded as outlined above.

### Pairwise sequence alignments

The C1^CARD^ (residues 1-96) and C9^CARD^ (residues 1-96) were subjected to a pairwise sequence alignment with the EMBOSS Needle webserver ^114^. This program performs a global sequence alignment using the Needleman-Wunsch algorithm^115^ with a default gap opening penalty of -10 and a gap extension penalty of -0.5. The alignment was scored using the BLOSUM62 substitution matrix. This resulted in an alignment with 26 matches and 12 gaps, corresponding to a pairwise sequence identity of 27.1%.

However, we observed that the default parameters in EMBOSS Needle yielded a relatively large number of gaps (12) relative to the lengths of the C1^CARD^ and C9^CARD^ sequences (96 residues). This resulted in an apparently suboptimal alignment, which contrasts with high structural similarity of the two domains (ca. 2-Å RMSD). Based on previous work by Elofsson ^116^, we modified the gap opening (-15) and gap extension penalties (-1), which were reported to produce better global alignments with the BLOSUM62 substitution matrix. These modifications introduced only two gaps in the alignment of C9^CARD^ and C1^CARD^ and produced a pairwise sequence identity of 19.8%. We implemented this in BioPython ^117^ version 1.85 and could produce highly similar alignments with the online EMBOSS Needle webserver when the corresponding parameters were updated (Gap Open = 15, End Gap Open = 15, End Gap = True, Gap Extend = 1, End Gap Extend = 1). Thus, we used these parameters in BioPython for the pairwise sequence alignment. For random sampling, we randomly selected *n* = 9 residues 10,000 times from the set of all C1^CARD^ surface residues (defined below) and computed the identity to C9^CARD^ at these sites based on the global alignment. The mean and standard deviation of the percent identity values were subsequently computed and compared to the observed value of 56%. A Z-score was calculated using the following equation:

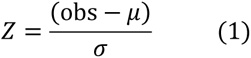

where *obs* refers to the observed pairwise sequence identity between the C9^CARD^ and C1^CARD^ at the type-1 filament interface (56%) and *μ* and *σ* to the mean and standard deviation of the sequence identities derived random sampling of surface residues as above (18 ± 12%).

### Relative solvent accessible surface area

We used DSSP^118^ version 4.4.7 to calculate the relative solvent accessible surface area (rSASA) for each residue in a PDB file. The BioPython ^117^ DSSP wrapper was used and its results parsed into an in-house Python script for downstream analysis. We supplied the PDB files 4rhw/chain E (C9^CARD^) and 5fna/chain A (C1^CARD^) for these calculations. Because the PDB file 5fna only contains structural data for residues 2-86 of the C1^CARD^, including an incomplete helix-6 that may produce non-realistic rSASA values, we also ran DSSP on the corresponding AlphaFold2^119^ structural model of the C1^CARD^ residues 1-96, which was obtained from the AlphaFold Database^120^ without further modification. The results presented include values derived from the AlphaFold2 model. To define surface from buried residues, we used an rSASA threshold of 25%, as reported previously^108^. The sequence identity of surface and buried residues was then determined from their positions in the global pairwise sequence alignment (as described above).

### ConSurf conservation scores

We extracted ConSurf conservation scores from the ConSurf-DB^121^ for PDB entries 4rhw, chain E (C9) and 5fna (C1). On the webserver, the reported grades (1-9) are derived from the raw ConSurf scores, with negative scores corresponding to more conserved positions. The assignment of grades can be extracted from the results summary (e.g. https://consurfdb.tau.ac.il/DB_NEW/4RHW/E/4RHW_E_consurf_grades.txt), and we report both values for ease of comparison to the webserver. Raw ConSurf scores for PDB 5fna are reported for residues 2-86, as these residues are present in the PDB file.

### NMR spectroscopy

All NMR samples were prepared in in 5-mm NMR tubes with 5% ^2^H_2_O added for the lock. The 2D ^1^H-^15^N HSQC spectrum in Figure 1 was recorded on a 23.5-T (1 GHz ^1^H frequency) Bruker Avance Neo spectrometer. To obtain resonance assignments of the C9^CARD^, a ^13^C,^15^N-labelled sample of C9^CARD^ was prepared at a protein concentration of 1.4 mM in 25 mM MES at pH 6.5 with 50 mM NaCl. Backbone (^1^H^N^, ^1^H^α^, ^15^N, ^13^CO, ^13^Cα) and side-chain (aliphatic ^13^C/^1^H) resonances were assigned based on three-dimensional (3D) HNCO, HN(CA)CO, HNCA, HN(CO)CA, HNCACB, C(CO)NH, H(CCO)NH spectra^122^ recorded at 20 °C on a 16.4-T (700 MHz ^1^H frequency) Bruker Avance-III spectrometer equipped with a cryogenically cooled, 5-mm TCI probe. Non-uniform sampling was employed in which the data sampling density in the indirect dimensions ranged from 10-25% and was based on a random sampling schedule. Spectral reconstruction was performed using SMILE^123^. All spectra were processed using NMRPipe^124^ and analyzed using NMRFAM-Sparky^125^. Secondary ^13^CA chemical shifts were determined by subtracting the neighbor-corrected random-coil chemical shifts predicted by POTENCI^126^. The 2D ^1^H-^15^N HSQC spectrum with a 100-ppm ^15^N sweep width was recorded on the same spectrometer, noting that the rectangular 180° pulses at this *B*_0_ on the ^15^N channel do not uniformly invert or refocus ^15^N spins that are located far off resonance^127^. However, given that we were simply interested in the chemical shifts of the aliased Arg side-chain signals – and not quantitative analysis of their intensities – we did not replace the rectangular pulses with adiabatic or other shaped pulses with broadband inversion profiles^127^.

^15^N longitudinal relaxation (*T*_1_), transverse rotating-frame relaxation (*T*_1ρ_)^128^ and CPMG relaxation dispersion experiments were collected on ^15^N-labeled C9^CARD^ at a protein concentration of 0.84 mM in 25 mM HEPES, 50 mM NaCl, 2 mM DTT at pH 6.5 with the temperature set to 25 °C. Data were recorded on a 14.1-T (600-MHz) Bruker Avance Neo NMR spectrometer with a room-temperature probe. The pulse sequences employed HSQC-based detection with sensitivity enhancement^129^. Magnetization was sampled at delay times of 0.01, 0.1, 0.25, 0.38, 0.5 (x2), 0.62, and 0.75 seconds, with a duplicated value at 0.5 seconds for error analysis. In the *T*_1ρ_ experiment, a 2-kHz ^15^N spin-lock field was applied to suppress chemical exchange during the transverse relaxation delay time (T), with ^1^H inversion pulses applied at T/4 and 3T/4 intervals to suppress cross-correlated relaxation between ^1^H-^15^N dipolar coupling and ^15^N chemical shift anisotropy ^130^ Magnetization was sampled at delay times of 5, 15, 30 (x2), 50, 75, 100, and 150 ms, with the 30-ms time point duplicated for error analysis. For the in-phase ^15^N-CPMG relaxation dispersion experiment^131^, 10 planes were collected with the constant-time relaxation delay set to 60 ms during which the ^15^N refocusing pulses were applied using a radiofrequency field strength of 5.8 kHz with [0013] phase cycling ^132,133^. For all three experiments described above, a total of 8 scans were collected per FID with an inter-scan delay of 3 seconds. The ^15^N and ^1^H spectral widths were 1,095 and 9,615 Hz with 50* and 615* complex points collected in each dimension, respectively corresponding to acquisition times of 45.6 and 64 ms. The ^15^N spin relaxation datasets were processed with NMRPipe, and NMRFAM-Sparky was used to pick peaks for subsequent lineshape fitting in FuDA^134^. The relaxation rates (i.e., 1/relaxation times) were determined by non-linear least-squares minimization of the *I/I_0_* intensity ratios to the function exp(-*R*t) where *R* is the fitted rate and *t* is the relaxation delay time. The ^15^N *R*_1ρ_ relaxation rates were corrected for the ^15^N spin-lock field strength and ^15^N resonance offset with the following equation:

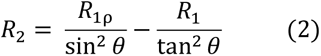

where *tan*θ = ω_1_/Ω in which ω_1_ is the spin-lock field strength and Ω the ^15^N resonance offset. The rotational correlation time was determined from the average ^15^N *T*_1_ and *T*_2_ relaxation times following the approach first outlined by Kay *et al.*^135^, with relaxation times included for residues located in regions of secondary structure. Values of the ^15^N effective transverse relaxation rate (*R*_2,eff_) were calculated using the following equation:

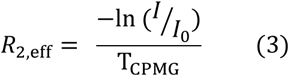

where T_CPMG_ is the constant-time relaxation delay, *I*_0_ is the peak intensity in the reference plane that lacks the constant-time delay, and *I* is the peak intensity in the other planes.

For the H38 p*K*_a_ determination, a 50 μM ^13^C,^15^N-labeled C9^CARD^ sample was prepared in 25 mM MES at with 50 mM NaCl buffer, and the pH was varied in 0.5-unit steps between 4 and 9.5 via the addition of concentrated HCl or NaOH aliquots. For each pH value, a 2D ^1^H-^13^C HSQC spectrum was recorded, and we focused on the His ^13^Cε chemical shift given that this site is sensitive to protonation of the imidazole ring but not impacted by the relative populations of the two different neutral tautomers (δ, ε) ^136–139^. The variation in the ^13^Cε chemical shift of H38 was fit to the Henderson-Hasselbalch equation as described^141^:

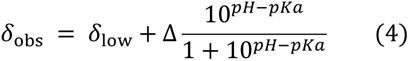

Where *δ*_obs_ is the observed chemical shift, *δ*_low_ the measured chemical shifts at pH 4, and Δ is the difference in chemical shift between the neutral and protonated forms. Least-squares fitting was performed in Python using the *lmfit* package ^140^, and the error was estimated from a standard bootstrap with 1,000 iterations.

### HydroNMR simulation

The C9^CARD^ crystal structure (4rhw, chain E) was supplied as input to the HydroNMR^82^ software program, which was used via the NMRbox resource^142^. In the simulation, the temperature was set to 25 °C to match the ^15^N relaxation experiments, with the solvent viscosity adjusted to 1 cPa. All other parameters, including the minimum and maximum radii of beads in the shell, were left at default values. The simulations assumed an N-H bond-length of 1.02 Å and a ^15^N chemical shift anisotropy of -160 ppm. The simulated rotational diffusion tensor of the C9^CARD^ yields values for *D*_xx_, *D*_yy_, and *D*_zz_ of 2.86, 2.89, and 3.39 x 10^7^ s^-1^, respectively, yielding an isotropic rotational correlation time of *D*_iso_ = 1/[6Tr(*D*)] = 5.47 ns, where Tr(*D*) corresponds to the trace of the 3x3 rotational diffusion matrix, or 1/3 (*D*_xx_ + *D*_yy_ + *D*_zz_). However, the rotational diffusion of C9^CARD^ appears approximately axially symmetric, with the anisotropic rotational diffusion values *D*_x_ = 2.90 x 10^7^ s^-1^, *D*_y_ = 2.79 x 10^7^ s^-1^, and *D*_z_ 3.43 x 10^7^ s^-1^ corresponding to values of 2*D*_z_ / (*D*_x_ + *D*_y_) = 1.20 and *D*_x_/*D*_y_ = 1.04.

### Differential scanning fluorimetry

To measure the melting temperature of C9^CARD^ variants, we mixed protein sample and SYPRO Orange dye (Thermo Fisher Scientific) in a 96-well plate (Bio-Rad) to a total volume of 25 µL per well. All conditions contained a protein concentration of 30 µM in 25 mM MES and 2 mM DTT at pH 5.5 or 7. SYPRO orange dye was added to each well in 1:2500 dilution, and the plate was then sealed. Fluorescence was measured in a Bio-Rad CFX96 RT-PCR instrument with the temperature increased from 20 °C to 95 °C in 0.5-°C increments every 30 seconds. The melting temperature was calculated from the first derivative of fluorescence versus temperature. All measurements were performed in triplicates, and the reported values are the mean and one standard deviation, normalized to the maximum and minimum values of each run.

### Determination of the maximum solubility limit of C9 variants

The maximum concentrations of soluble C9^CARD^ and related C9 variants were determined at 25 °C at pH 5.5 or pH 7 (25 mM MES, 2 mM DTT, 0.5 mM EDTA, 0.04% (w/v) NaN_3_). Purified protein was concentrated in an Amicon 3-kDa molecular weight cutoff spin filter (Amicon^®^ Ultra-15 centrifugal filter) at 3.5k x *g* with the temperature in the centrifuge (Eppendorf centrifuge 5810 R) pre-equilibrated to 25 °C. The sample was then transferred to a 1.5-mL Eppendorf tube and further centrifuged at 20k x *g* for 2 minutes at 25 °C to pellet the remaining insoluble material. The concentration of the supernatant (*i.e.*, *C*_sat_) was then determined using a Nanodrop (Thermo Scientific NanoDrop One) via aborbance at 280 nm and the corresponding extinction coefficient. This was performed in triplicate for each C9 variant at each pH, and the mean and standard deviation *C*_sat_ values are reported.

### Negative-stain electron microscopy

A carbon-coated copper grid (400-mesh) (Science Services, Germany) was glow-discharged using a PELCO easiGlow™ Glow Discharge Cleaning System (Ted Pella Inc., USA)) to make it hydrophilic for homogenous sample distribution. 10 μL of purified C9^CARD^ (25 mM HEPES, 200 mM NaCl, 0.5 mM EDTA, 2 mM TCEP, pH 7.5) was adsorbed onto the glow-discharged grid and allowed to settle for 1 minute. The remaining sample was carefully removed with filter paper, and immediately 10 μL of a 1% (w/v) uranyl acetate staining solution was added. The uranyl acetate solution was allowed to settle for 1 minute and the excess solution was removed using filter paper. The sample was air dried for few minutes before putting inside the microscope. Sample visualization was done on a Tecnai G2 FEI (Thermo Fisher Scientific Inc., USA) microscope equipped with an Ultrascan 1000 CCD camera (Gatan AMETEK, Germany) operating at an acceleration voltage of 120 kV.

### Cryo-electron microscopy

For the cryo-EM structural analyses, we induced WT and H38R C9^CARD^ filament formation by concentrating the purified proteins at 25 °C to the desired C*_sat_*, which resulted in long and straight filament bundles (**Supplementary Figure 7, Supplementary Figure 13**). The grids were prepared and vitrified at the Cryo-Electron Microscopy Platform (CEMP) of Helmholtz Munich and the IBS, Grenoble EM facility. For cryo-EM grid preparation, 4.5 µL of the C9^CARD^ sample was applied to a glow-discharged 200 mesh Quantifoil R2/1 grid, blotted for 4 s with force 4 in a Vitrobot Mark III (Thermo Fisher) at 100% humidity and 4 °C. Grids were plunge frozen in liquid ethane-propane mixture, cooled by liquid nitrogen. Cryo-EM data acquisition was performed with a 300-kV, FEI Titan Krios transmission electron microscope (Thermo Fisher) with EPU software. Movie frames were recorded at a magnification of 130,000x using a Falcon4i direct electron detector camera. For the H38R C9^CARD^ filament, the total electron dose of about 55 electrons per Å^2^ was distributed over 30 frames at two different pixel sizes of 0.76 Å and 0.95 Å. The two collected datasets were first processed separately, and the final particle sets were then merged by rescaling the particles from 0.76 Å dataset to a pixel size of 0.95 Å. For the WT C9^CARD^ filament sample, data were recorded at a single pixel size of 0.76 Å. Movies were recorded with a defocus range of −0.75 to −2.5 μm. Data processing was performed using the CryoSparc v4.4.1 helical reconstruction pipeline^112^ (**Supplementary Figure 7, Supplementary Figure 13**). The picking of filaments was done in several steps. First around 200 segments were selected manually, and the resulting best 2D class-average was used for template-based filament tracing. Subsequently, the best 2D class-averages were used for a new filament tracer job, and a final re-picking of the dataset was done using as templates the 2D projections of the first obtained 3D reconstructions. In all cases, the segments were extracted with a separation distance of 27 Å in a box of 340 Å. H38R C9^CARD^ filament helical parameters were estimated by indexing the layer lines of the sum of the power spectra of the segments belonging to the most detailed 2D class using Helixplorer (http://rico.ibs.fr/helixplorer/). However, for the WT C9^CARD^ filament, we used the symmetry parameters from the H38R C9^CARD^ filament as a starting point for refinement. Using the estimated symmetry parameters based on Helixplorer, homogenous refinement was performed with 537,539 and 873,929 particles corresponding to H38R and WT C9^CARD^ filaments respectively. After multiple rounds of 3D classification, 291,659 and 361,284 segments of H38R and WT dataset were selected for the final refinement. Particles selected after 3D classification were re-extracted without binning and used for 3D non-uniform (NU) refinement. A final round of NU-refinement followed by CTF refinement led to a refined helical symmetry for H38R C9^CARD^ (H38R)- rise: 9.13 Å, pitch: 48.64 Å, twist: -67.2° and WT C9^CARD^- rise: 9.27 Å, pitch: 49.04 Å, and twist: -68.04°.

The half-maps for the H38R and WT C9^CARD^ filaments were imported in RELION^143^ for post-processing. The estimated resolution based on Gold Standard Fourier Shell Correlation (GS-FSC) 0.143 criteria, and by applying a b-factor of -90 Å^2^ was 3.5 Å and 3.3 Å for H38R and WT C9^CARD^ filaments respectively (**Supplementary Figure 7, Supplementary Figure 13**). For both the structures, to visualize all three interfaces, a filament model containing seven subunits was fitted via rigid body fitting of the C9^CARD^ crystal structure (3YGS; chain P) into the obtained cryo-EM density maps. Manual adjustments were performed in Coot^144^, followed by real-space refinement using Phenix 1.20.1^146^ to obtain the final WT and H38R C9^CARD^ filament models. The map versus model FSC (**Supplementary Figure 7, Supplementary Figure 13**) was calculated by Phenix ^146^. The final structure consists of residues 1 to 99 with a preceding Ser-Gly overhang. Interfacial residues and surface area calculations were performed using PDBePISA^145^ (https://www.ebi.ac.uk/pdbe/pisa/).

### Structure-based pK_a_ prediction

C9^CARD^ PDB files (PDB ID/chain: 4rhw/E, 3ygs/P, 5wvc/B, AlphaFoldDB/A, 5juy/O, 5wve/S) were uploaded to the PDB2PQR webserver that assigns titration states with version 3.6.1 of PROPKA^89^. The pH of the calculation was set to 7.0 and the default forcefield was used. Steric clashes were removed with the debump algorithm inside PDB2PQR that optimizes inter-atomic distances that fall below defined threshold values of 1.0 Å for hydrogen-hydrogen, 1.5 Å for hydrogen-heavy atom, and 2.0 Å for heavy atom-heavy atom collisions. Hydrogen bond networks were optimized by PDB2PQR by enabling the flipping of His, Asn, and Gln side chains, the rotation of side-chain hydrogen atoms on Ser, Thr, Tyr, and Cys residues, and placement of side-chain hydrogen atoms on His, Asp, and Glu.

### Molecular dynamics simulations

We performed all-atom molecular dynamics simulations with the GROMACS^147^ simulation package using the AMBER99SB-ILDN force field^148^. We selected the highest-resolution crystal structure of the C9^CARD^ (PDB: 4rhw) and extracted chain E from the PDB file. The system was solvated in a rhombic dodecahedron of TIP4P-EW^149^ water molecules with a minimal distance of 10 Å between the protein and the box edge, and then energy minimized using the steepest descent algorithm to remove any steric clashes, with the maximum force set to 1000 kJ mol^-1^ nm^-1^. Following minimization, the system was equilibrated in two steps: *NVT* equilibration at 300 K for 100 ps, using the v-rescale thermostat^150^, and *NPT* equilibration at 300 K and 1 bar for an additional 100 ps, with Berendsen pressure-coupling^151^. During equilibration, all protein heavy atoms were restrained to their initial positions. Production MD simulations were carried out in the *NPT* ensemble at 300 K and 1 bar using periodic boundary conditions. The particle mesh Ewald method^152^ was employed for long-range electrostatic interactions beyond 1.0 nm, and we used a 1.0 nm cutoff for van der Waals interactions. The simulation time step was set to 2 fs, and coordinates were saved every 10 ps. The temperature was maintained at 300 K using the v-rescale thermostat, while pressure was controlled using the Parrinello-Rahman barostat^153^. All simulations were performed on a single node utilizing 32 CPUs or one GPU. For each system, we discarded the first 50-ns of production MD simulations and quantitatively analyzed the subsequent 5 μs. For the neutral state of H38, the proton was placed on the ND1 position. For protonated H38, we placed protons at both the ND1 and NE1 positions, creating the cationic form or imidazolium. For the H38R variant, we performed *in silico* mutagenesis in PyMOL to introduce the R38 residue. Overall charge neutrality was maintained in all simulations through the addition of counter ions.

Analysis of the trajectories was performed with GROMACS scripts and the MDTraj^154^ python package. Side-chain χ_N_ torsion angles were computed with the in-built *compute_chiN* functions in MDTraj where *N* = 1, 2, 3, or 4. We defined *gauche*+, *trans*, and *gauche*- rotamers as angles that fall within ±20° of 60°, 180°, and 300°, respectively^99^. To determine the populations of different χ_1_ rotameric states for eachm residue (except for Ala and Gly), we plotted a histogram of χ_1_ torsion angles (bin size = 2 °) over the course of the trajectory and integrated the bins over the ranges defined above. The populations of each state were then normalized such that the sum was unity for each residue. For the 1D histograms shown in Figure 3, the distributions were smoothed with a Savitzky-Golay filter with filter window length of 15 and polynomial order 3.

### Structural bioinformatics

Helical protein filaments were downloaded from the PDB in February 2025 using the following search criteria: EM Reconstruction Method (“Helical”), Reconstruction Resolution (<= 5.0 Angstroms), Polymer Entity Sequence Length (“>= 50 residues”), PubMed Abstract (has any of the words: “filament”), and PubMed Abstract (does not have any of the words: (“amyloid”, “tauopathies”, “amyotrophic”, “Parkinson’s”, “Alzheimer’s”, “dementia”, “frontotemporal”). The latter query was included to remove amyloid fibrils whose structures were determined by helical reconstruction: e.g., there are 160 amyloid fibril structures (tau, SOD1, alpha-synuclein, TAF-15, etc.) returned from the above search when amyloid-related terms are included. Overall, this query yielded 173 PDB files that were included for downstream analysis. To reduce sequence redundancy, we clustered the sequences with CD-HIT^155^ to maximum sequence identities of 90%, 70%, and 50%, respectively yielding 77, 70, and 65 clusters.

### AF2 ECOD

We downloaded the AF2 Evolutionary Classification Of Domains (ECOD)^102^ on 3 January 2025 via the link: http://prodata.swmed.edu/ecod/human/distribution. For each AlphaFold2-predicted domain structure, we ran DSSP as described above (see *Relative solvent accessible surface area*) to assign per-residue secondary structure types. We then collected all helical residues to quantify and compare the residue types at each helical position. We used the Richardson and Richardson terminology^95^ to define helical positions: N-cap, N1, N2, N3, [middle], C3, C2, C1, C-cap define a helix, with [middle] corresponding to the intervening or middle helical residues and N1-N3 (C3-C1) the first (final) three residues with helical torsion angles that contribute only carbonyl (amide) groups to the backbone hydrogen bond network. The N-cap and C-cap residues are the preceding and following residues, respectively, that generally form hydrogen bonds with the terminal backbone or side-chain residues. We then extracted residue types at each position across all helices in the DSSP-mapped AF2 ECOD dataset. To quantify the fold enrichment of D and R residues in the N1/C1, N2/C2, and N3/C3 positions, we first defined the background D and R frequencies across all helical residues, regardless of residue position. The frequency of a given residue type (X) in

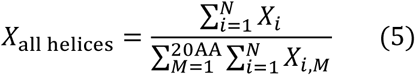

Where *i* sums over all helices in the dataset and *M* denotes all 20 residue types. The fold enrichment at a given helical residue position is then:

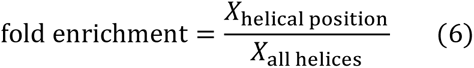

where *X*_helical position_ is defined in the same manner as *X*_all helices_, except that the sums in the numerator and denominator reflect the number of *X* residues observed at a particular helical position (as opposed to in all helical residues). For example, for *X*_N3_ with X = R or D, there are 73,140 total N3 residues of which 2,795 are R (0.038) and 5,243 are D (0.072). In all helices, for *X*_all helices_ with X = R or D, there are 1,764,346 residues of which 105,904 are R (0.060) and 62,107 are D (0.035). The fold enrichments for R and D at the N3 position are therefore 0.63 (0.038/0.060) and 2.06 (0.072/0.035), respectively. For the correlation plots by helical residue position, we imposed a minimum helix length of 12 residues to enable sufficient distance between the helical termini and middle residues, defined here as the N5 and C5 positions (i.e. N-cap, N1, N2, N3, N4, N5, C5, C4, C3, C2, C1, C-cap corresponds to a 10-residue helix with N5 and C5 being the 5^th^ and 6^th^ helical residues).

### Human Domainome 1

Human Domainome 1 is a previously established resource from the Lehner laboratory^108^ and contains 522 protein domains that were subjected to single-site saturation mutagenesis. There are 602,882 total mutations in this dataset. The fitness of each single-site variant was obtained through a fragment complementation assay that couples variant abundance with cell growth. This parameter correlates with thermodynamic stability. We ran DSSP as described above (see *Relative solvent accessible surface area)* on the AlphaFold2-predicted structures of each domain to identify secondary structure. All DSSP-defined helical residues were then extracted, including the preceding and following residues (N- and C-cap, respectively) and then used to extract the corresponding mutations in the Human Domainome 1 dataset. We considered helices of seven residues or longer, to exclude short helices that contain fewer than two turns. Thus, we could compare use this filtered dataset to then assess the impact on protein fitness when introducing an R vs. D residue across many different protein domains, as we had performed with our H38R vs. H38D mutations in the C9^CARD^.

For each helix, we extracted X-to-R and X-to-D mutations, where X is defined as any polar residue, with X = [A, C, G, H, M, N, P, Q, S, T] to exclude sites that natively contain charged or bulky hydrophobic residues that may be less tolerant to mutations. We then determined the helix-normalized fitness defect at each position in each helix by calculating the fitness difference between the X-to-R/D mutation and the median fitness of all mutations to polar residues at this site (i.e. fitness(X-to-R/D) – median(fitness(X-to-Y)), where Y exists in X but does not equal X). We excluded P from the definition of Y, as P residues are generally highly destabilizing across the Human Domainome 1 dataset^108^, especially in regions of secondary structure such as the middle of helices. Values that are greater than zero, therefore, reflect X-to-R/D mutations that have higher fitness (i.e., more thermodynamically stable). Finally, we then separated our helical residues by their relative position within each helix: N-cap, N1, N2, and N3 comprise the N-helix residues; C-cap, C1, C2, and C3 comprise the C-helix residues; and residues in between are defined as the helix middle.

For the two CARD domains within the Human Domainome 1 dataset (RIPK2^CARD^, ASC^CARD^), we identified N-cap, N1, N2, or N3 residues that are negatively charged or polar. We then collected the normalized fitness values in the Human Domainome 1 for the following mutation types: WT to negative (E/D-to-D/E or polar-to-E/D), WT to positive (E/D-to-R/K or polar-to-R/K), WT to stop codon (E/D-to-* or polar-to-*), and the three most tolerated mutations excluding charged residues. The latter category provides an estimate of the intrinsic sensitivity to mutations at that particular position, i.e. the degree to which conservative mutations can be tolerated. Nonsense mutations, on the other hand, provide a baseline of the minimum normalized fitness level that is expected for a truncated and misfolded protein.

## Acknowledgements

We thank Lewis E. Kay (University of Toronto), Michael Sattler (Technical University of Munich, Helmholtz Center Munich), and thesis committee member Malene Ringkjobing Jensen (IBS Grenoble) for their support, feedback, and insightful discussions. Christian Loew and Siavash Mostafavi are acknowledged for their advice with cryo-EM sample preparation. This study made use of NMRbox: National Center for Biomolecular NMR Data Processing and Analysis, a Biomedical Technology Research Resource (BTRR), which is supported by NIH grant P41GM111135 (NIGMS). S.R. was trained within the framework of the PhD program BioMolStruct. T.M. is grateful to the Austrian Science Fund (FWF) for excellence cluster 10.55776/COE14, Grants DOI 10.55776/P28854, 10.55776/I3792, 10.55776/DOC130, and 10.55776/W1226, the Austrian Research Promotion Agency (FFG) grants 864690 and 870454; the Integrative Metabolism Research Center Graz; the Austrian Infrastructure Program 2016/2017; the Styrian Government (Zukunftsfonds, doc.fund program); the City of Graz; and BioTechMed-Graz (flagship project). This project was funded in part by the FFG and the European Union (EFRE) under grant 912192. S.R. and T.M. acknowledge funding from the Medical University of Graz (MUG Cryo-EM funding), the Center for Medical Research for laboratory access, and thank the Vienna BioCenter cryo-EM facility and Thomas Heuser for the initial screening. S.R. acknowledges funding from the MUG mobility grant for female scientists and the Marietta-Blau Stipendium from the Austrian Agency for Education and Internationalization (OEAD). S.R. and D.K. thank Dominique Pernitsch for the help with negative staining TEM. A.D. and S.R. thank MICA team. S.R. and A.D. thank Guy Schoehn for establishing and managing the IBS cryo-EM platform and for providing training and support, and Lefteris Zarkadas for assistance at the Glacios microscope. We used ChatGPT from OpenAI to aid in editing grammar and written style. We used the platforms of the Grenoble Instruct-ERIC centre (ISBG; UAR 3518 CNRS-CEA-UGA-EMBL) within the Grenoble Partnership for Structural Biology (PSB), supported by FRISBI (ANR-10-INBS-0005-02) and GRAL, financed within the University Grenoble Alpes graduate school (Ecoles Universitaires de Recherche) CBH-EUR-GS (ANR-17-EURE-0003). The IBS EM facility is supported by the Rhône-Alpes Region, the Fondation Recherche Medicale (FRM), the fonds FEDER and the GIS-Infrastrutures en Biologie Sante et Agronomie (IBISA). We thank the Helmholtz Munich Digital Transformation and IT Department (DigIT) team for support of computational resources. T.R.A. acknowledges funding from the Initiative and Networking Fund of the Helmholtz Association (Helmholtz Investigator Grant VH-NG-20-14).

## Author Contributions

T.R.A. conceived and designed the study. S.R. and T.R.A. expressed and purified protein samples. S.R. performed DSF thermal melts and analyzed them with T.R.A. S.R., B.B., T.M., and T.R.A collected NMR data. T.R.A processed and assigned the 3D NMR data. T.R.A processed and analyzed the NMR spin-relaxation datasets. S.R. performed the saturation concentration experiments. S.R., B.B., and T.M. performed and processed the p*K*_a_ measurement, which was fit by T.R.A. S.R optimized the EM samples. S.R., M.B., and D.K. collected negative-stain data. S.R., M.B., and S.B. collected cryo-EM data. S.R., S.B., T. P.-K., and A.D. analyzed EM data, which was processed and analyzed by S.R. and A.D. T.R.A. produced and analyzed the molecular dynamics simulations. Ð.K., I.P., and T.R.A. performed the pairwise sequence alignments and random sampling approach on C1 and C9 paralogs. T.R.A. performed the structural bioinformatics experiments on the AF2 ECOD and Human Domainome 1 datasets with input from I.P. T.R.A. and S.R. prepared the figures. T.R.A. wrote the initial manuscript with S.R., and all authors commented and contributed to the approved final version of the manuscript.

## CRediT format

**Swasti Rawal** – Investigation, Data Curation, Formal Analysis, Project Administration, Writing – Initial Draft, Writing – Reviewing & Editing, Visualization, Funding Acquisition. **Stefan Bohn** – Investigation, Data Curation, Writing – Reviewing & Editing. **Maria Bacia** – Investigation, Formal Analysis, Writing – Reviewing and Editing. **Ðesika Kolarić** – Investigation, Data Curation, Formal Analysis, Writing – Reviewing & Editing. **Benjamin Bourgeois** – Investigation, Formal Analysis, Writing – Reviewing & Editing. **Dagmar Kolb** – Investigation, Writing – Reviewing & Editing. **Tea Pavkov-Keller** – Formal Analysis, Supervision, Resources, Writing – Reviewing & Editing. **Iva Pritišanac** – Investigation, Formal Analysis, Resources, Writing – Reviewing & Editing. **Tobias Madl** – Data Curation, Formal Analysis, Resources, Supervision, Funding Acquisition, Writing – Reviewing & Editing. **Ambroise Desfosses** – Investigation, Resources, Data Curation, Formal Analysis, Methodology, Supervision, Writing – Reviewing & Editing. **T. Reid Alderson** – Conceptualization, Investigation, Data Curation, Formal Analysis, Resources, Supervision, Project Administration, Writing – Original Draft, Writing Reviewing & Editing, Visualization, Funding Acquisition.

