## Supplementary Information for "A histidine switch controls the pH-responsive self-assembly of a helical protein filament"

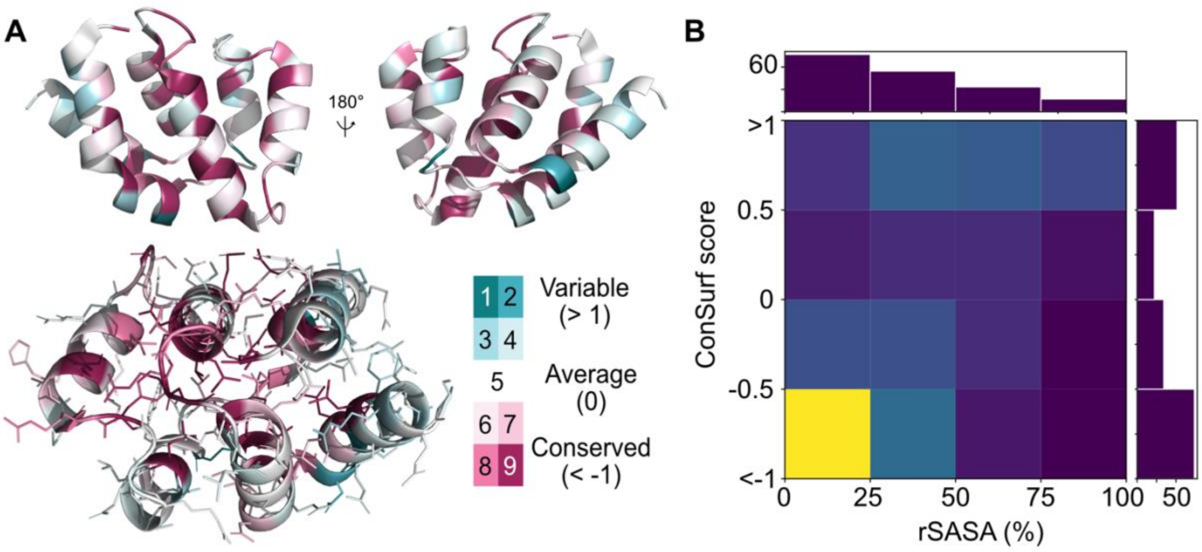


**Supplementary Figure 1. Comparison of C9$^{CARD}$ and C1$^{CARD}$ sequences.**
(**A**) ConSurfDB[65]-derived sequence identity among C9 orthologs mapped onto the structure of the C9$^{CARD}$ (PDB: 4rhw). The degree of conservation is shown with the color bar. Amino-acid side chains show that buried residues appear more conserved than those on the surface. (**B**) 2D histogram showing the per-residue relative solvent accessible surface area (rSASA) versus the ConSurf score for both the C9$^{CARD}$ and C1$^{CARD}$. The number of residues in each column from 0-25, 25-50, 50-75, and 75-100% rSASA is $n$ = 76, 54, 33, and 17, respectively.

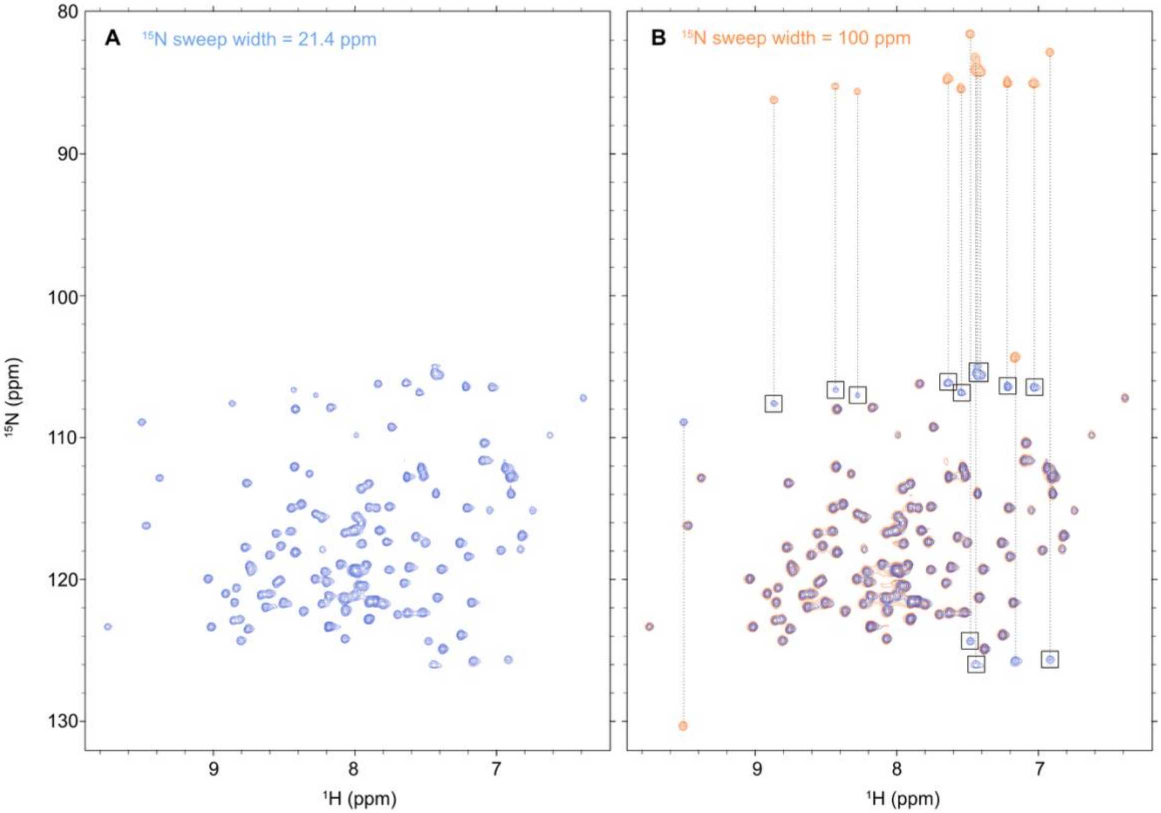


**Supplementary Figure 2. Additional signals from Arg Nε-Hε groups in the 2D $^{1}$H-$^{15}$N HSQC spectrum of C9$^{CARD}$.** Comparison of 2D $^{1}$H-$^{15}$N HSQC spectra recorded on the same sample of $^{15}$N-labeled C9$^{CARD}$ at 25 °C in pH 6.5 buffer (25 mM MES, 50 mM NaCl) at 700 MHz $^{1}$H Larmor frequency with the $^{15}$N sweep width set to (**A**) 21.4 ppm or (**B**) 100 ppm. The $^{15}$N carrier frequency was the same in both experiments (115.45 ppm), and the spectrum was set up such that aliased peaks did not acquire a 180° phase shift. In panel **B**, the two spectra are overlaid (blue, orange), and vertical dashed lines connect the aliased peaks (blue) with the peaks that have their true $^{15}$N chemical shifts (orange). Rectangles are drawn around the blue peaks that originate from Arg Nε-Hε side-chain groups, which resonate near 85 ppm in the $^{15}$N dimension and are thus aliased in panel A.

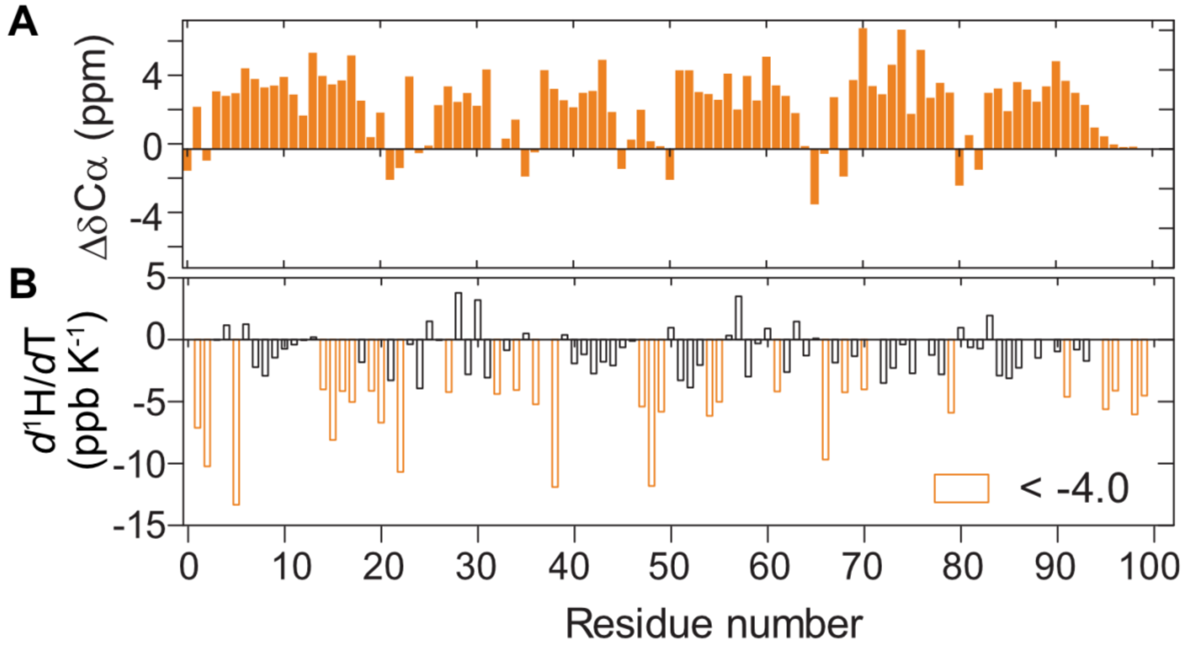


**Supplementary Figure 3.** (**A**) Secondary $^{13}$Cα chemical shifts (Δδ$^{13}$Cα) and (**B**) $^{1}H^{N}$ temperature coefficients ($d^{1}H/dT$) for the C9$^{CARD}$. Residues with $d^{1}H/dT$ values below -4.0 ppb/K, suggestive of non-hydrogen-bonded amides, are shown in orange.

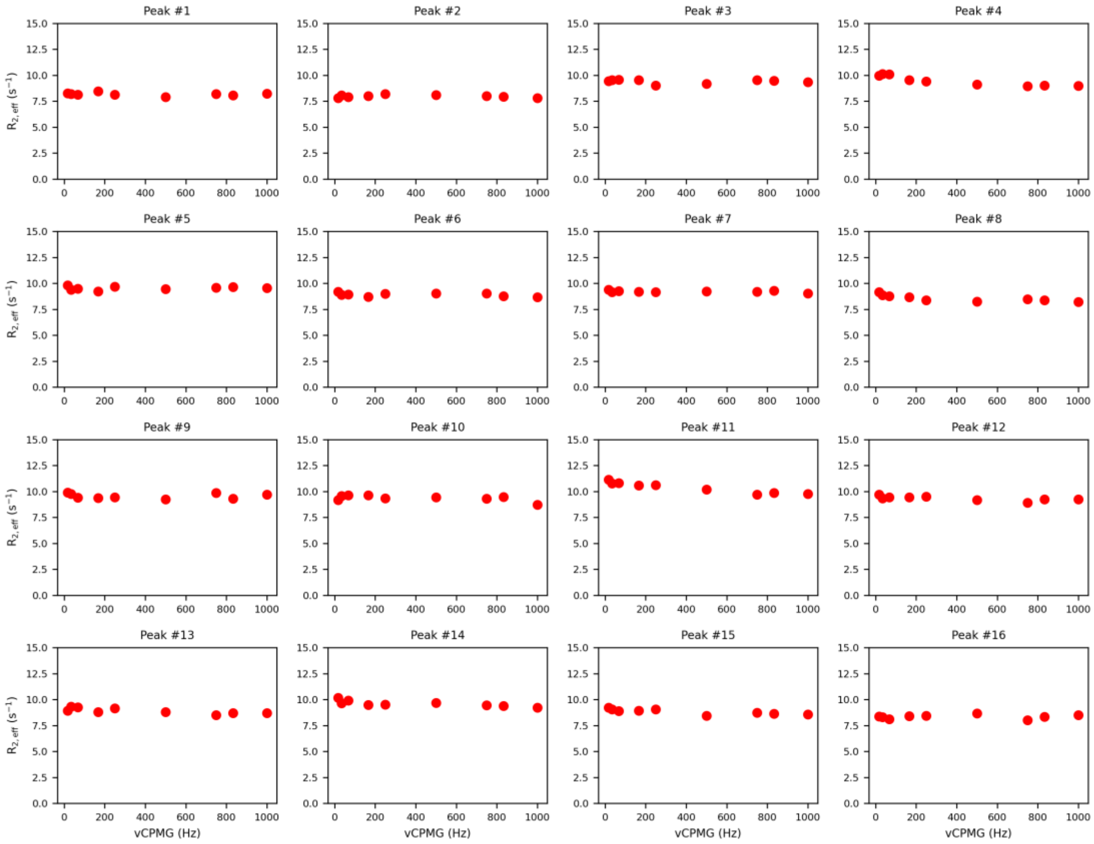
Peak #1
Peak #2
Peak #3
Peak #4
Peak #5
Peak #6
Peak #7
Peak #8
Peak #9
Peak #10
Peak #11
Peak #12
Peak #13
Peak #14
Peak #15
Peak #16
$R_{2,\,eff}$ ($s^{-1}$)
νCPMG (Hz)

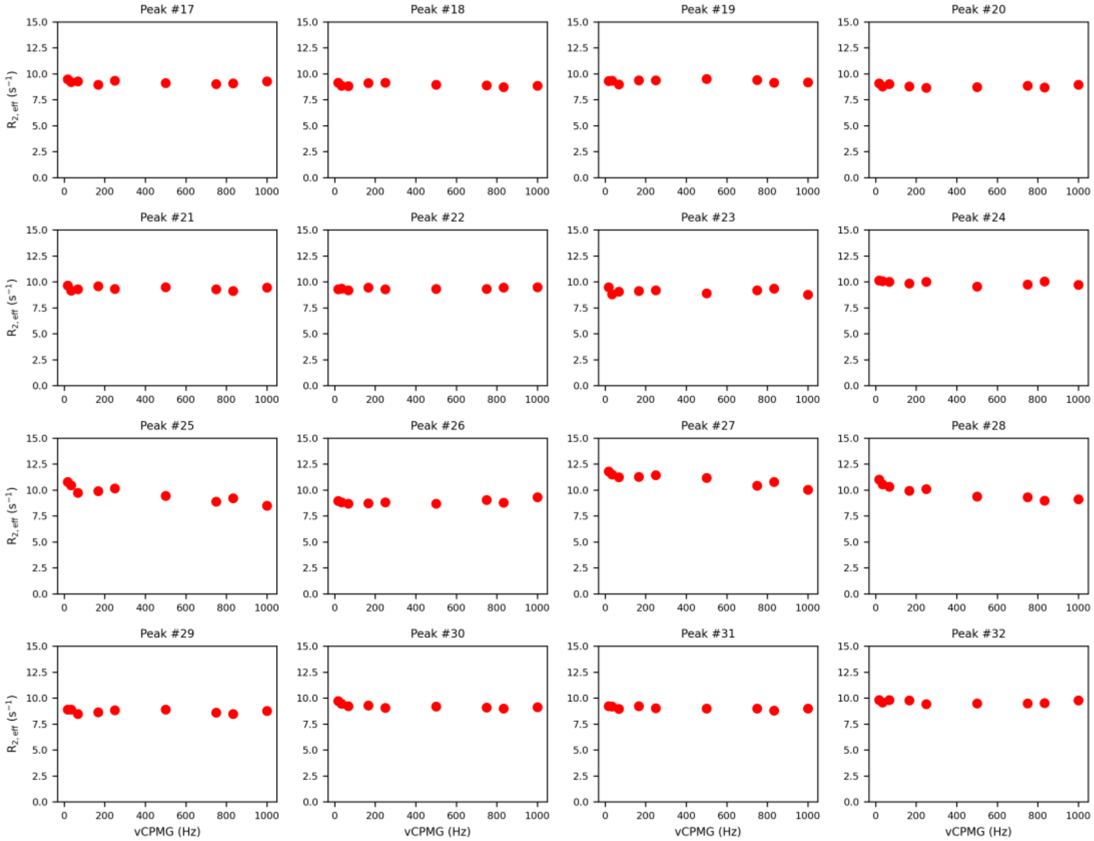
Peak #17
Peak #18
Peak #19
Peak #20
Peak #21
Peak #22
Peak #23
Peak #24
Peak #25
Peak #26
Peak #27
Peak #28
Peak #29
Peak #30
Peak #31
Peak #32
$R_{2,\,eff}$ ($s^{-1}$)
νCPMG (Hz)

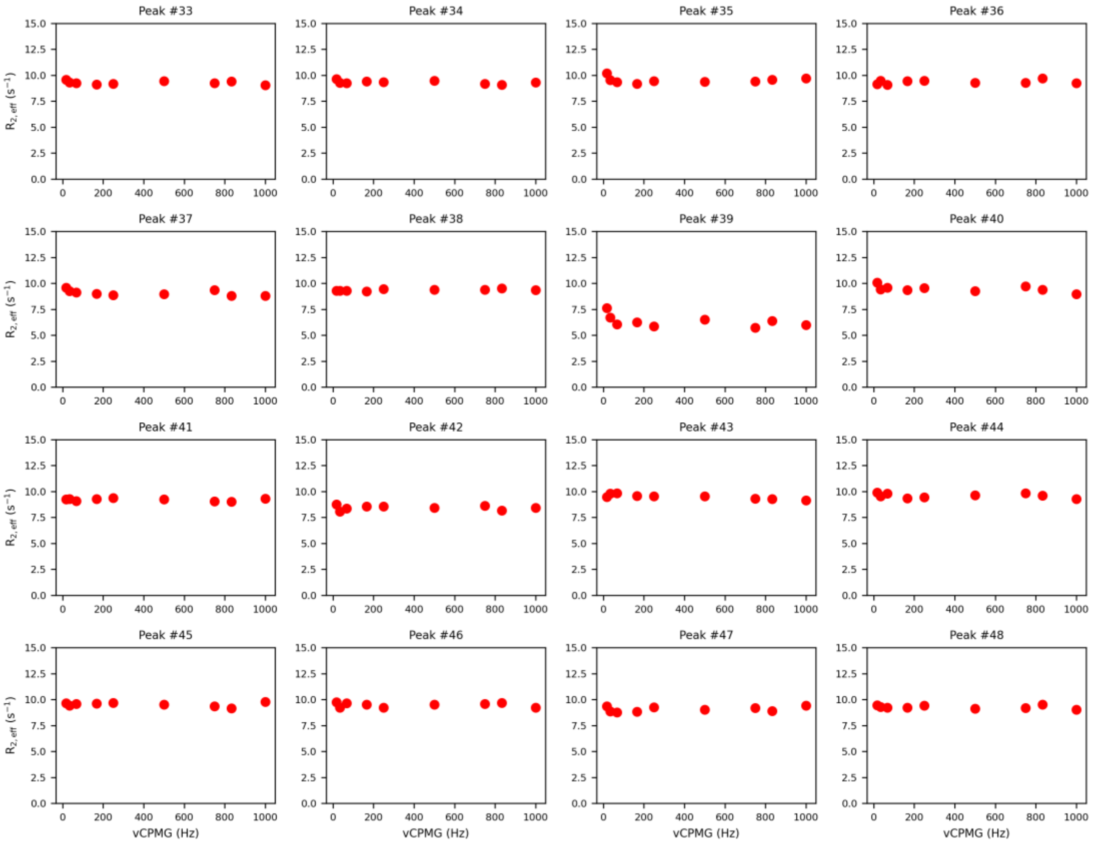

Peak #33
Peak #34
Peak #35
Peak #36
Peak #37
Peak #38
Peak #39
Peak #40
Peak #41
Peak #42
Peak #43
Peak #44
Peak #45
Peak #46
Peak #47
Peak #48
$R_{2,eff}$ ($s^{-1}$)
vCPMG (Hz)


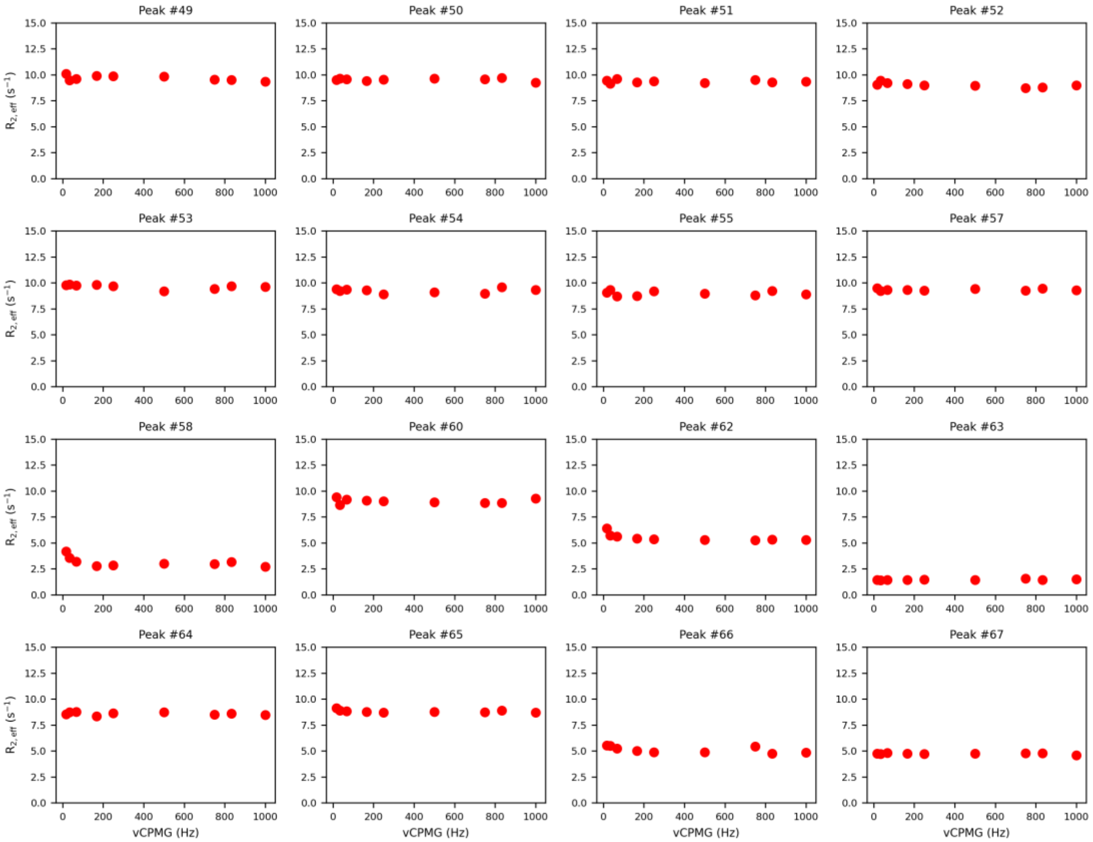

Peak #49
Peak #50
Peak #51
Peak #52
Peak #53
Peak #54
Peak #55
Peak #57
Peak #58
Peak #60
Peak #62
Peak #63
Peak #64
Peak #65
Peak #66
Peak #67
$R_{2,eff}$ ($s^{-1}$)
vCPMG (Hz)

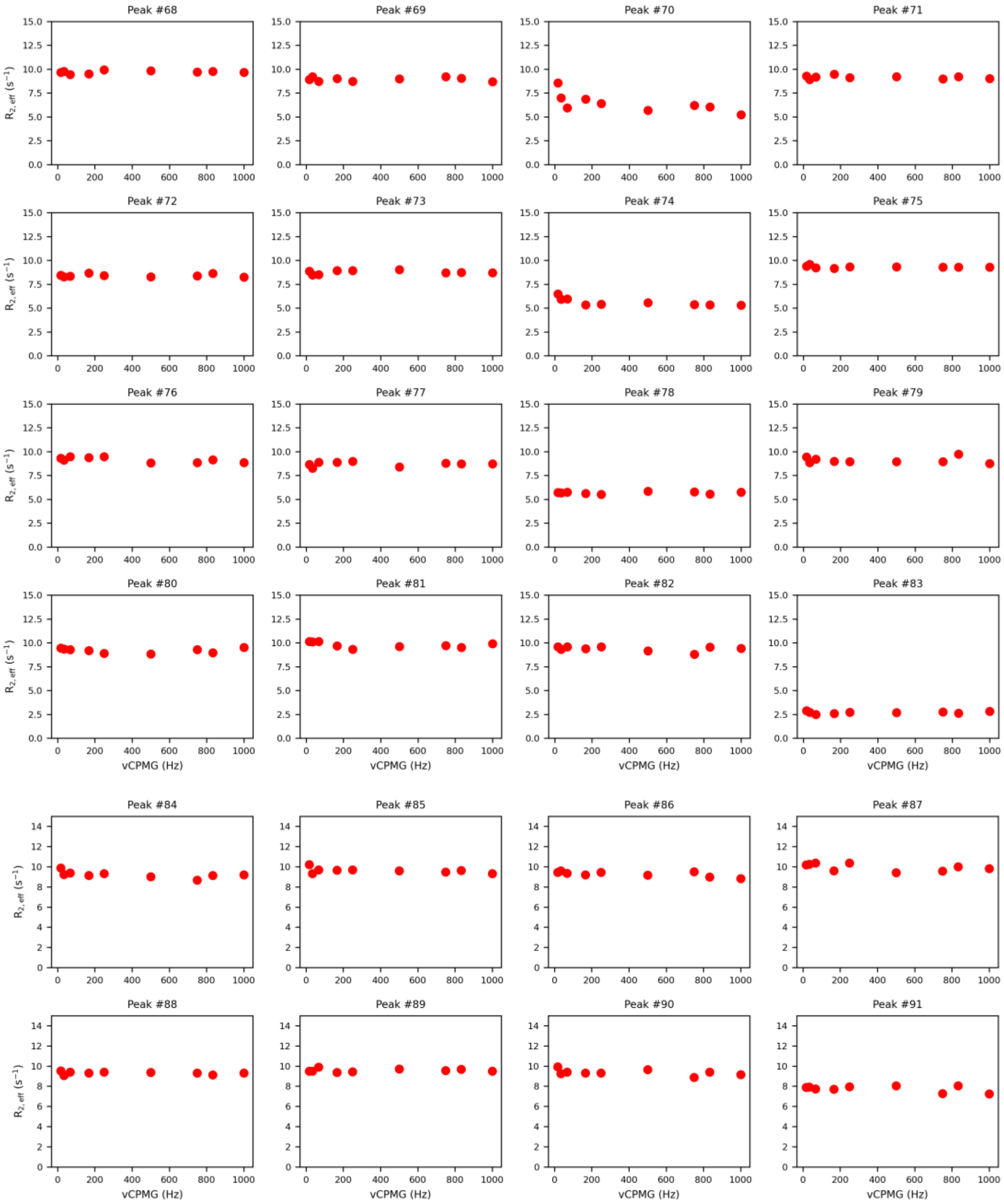


**Supplementary Figure 4. $^{15}$N-CPMG relaxation dispersion does not show evidence of transient oligomerization on the millisecond timescale.** CPMG relaxation dispersion data were collected on a $^{15}$N-labeled sample of C9$^{CARD}$ at a protein concentration of 0.84 mM in 25 mM HEPES, 50 mM NaCl, 1 mM EDTA at pH 6.5 and 298 K.

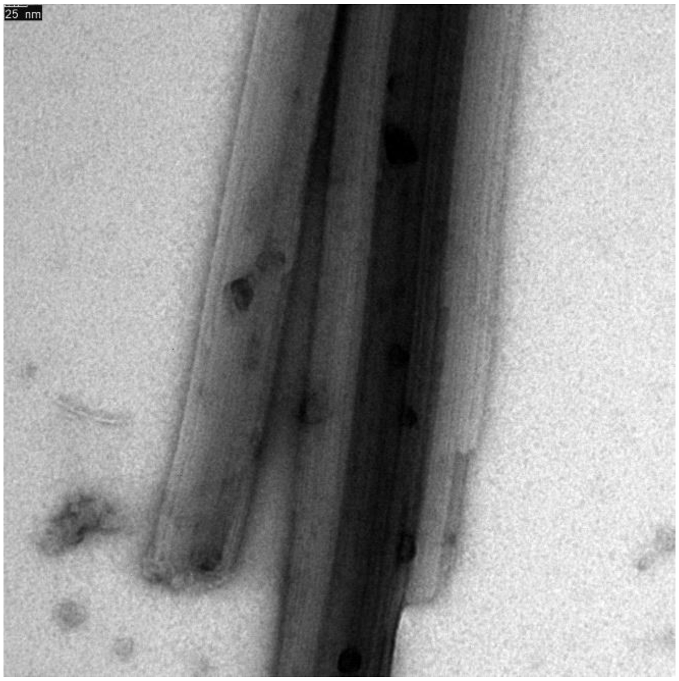


**Supplementary Figure 5. The aggregates of the CARD formed at neutral pH are also filamentous.**

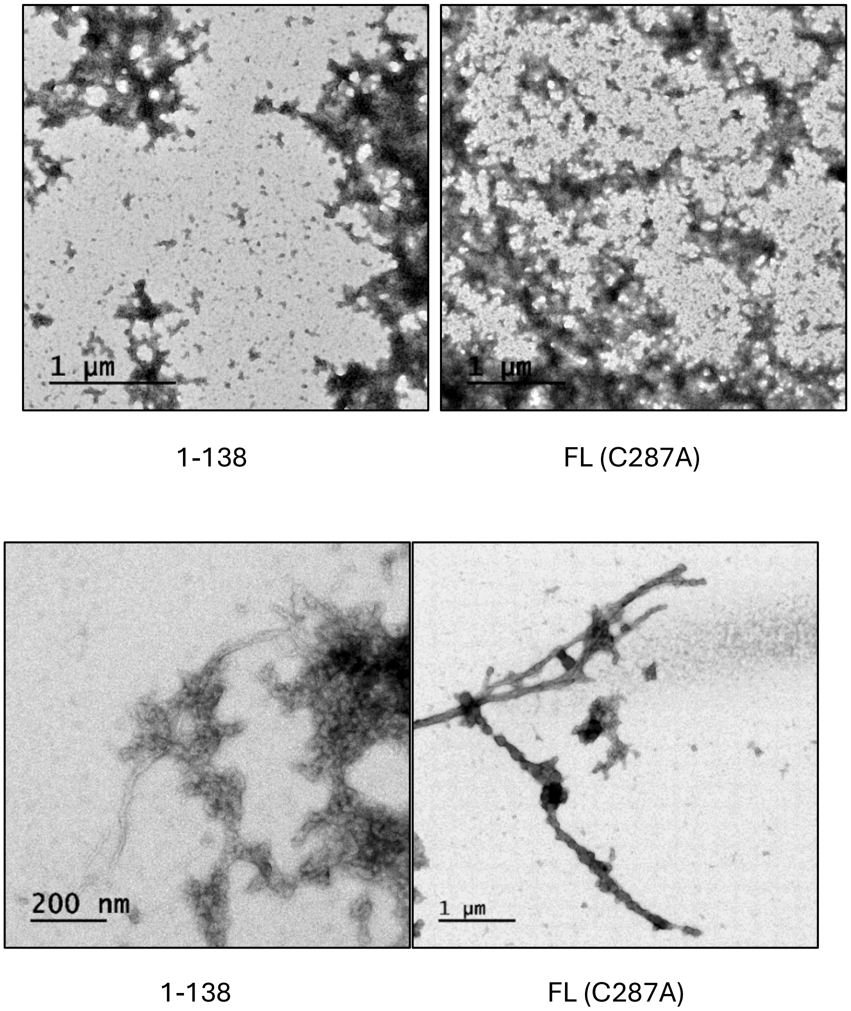


**Supplementary Figure 6**. Negative-stain EM images of C9 CARD+linker (1-138) and full-length C9 with the C287A mutation. The upper row shows representative micrographs in which amorphous aggregates are observed, which are abundant. In a few areas, filaments as observed in the bottom row, are observed. However, filaments represent a small minority of the grid.

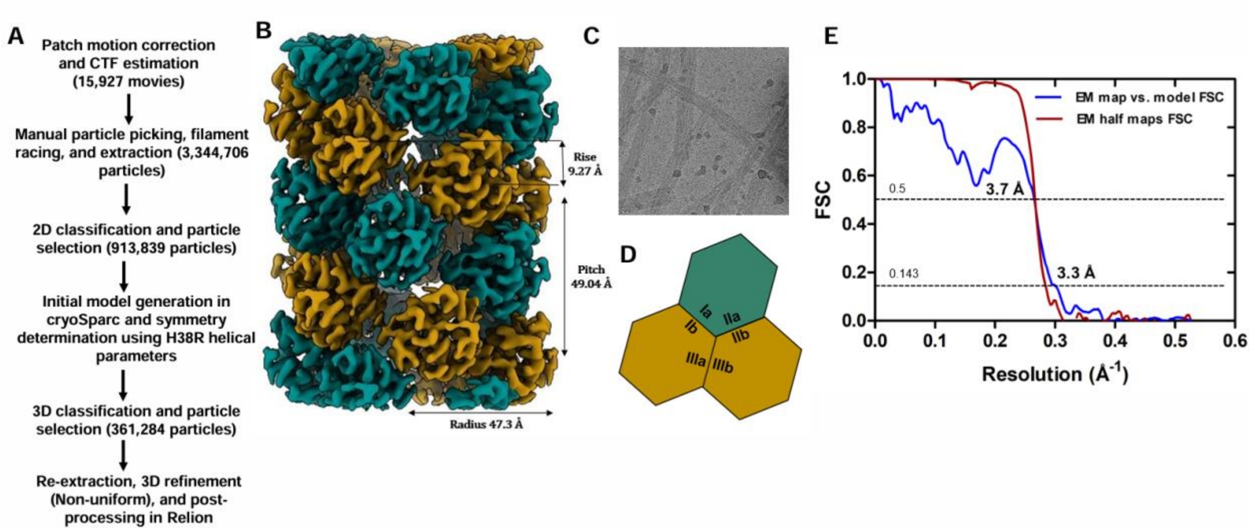


**Supplementary Figure 7. Structure determination of WT C9$^{CARD}$ filament** (**A**) Data processing pipeline for 3D reconstruction using CryoSparc and Relion. (**B**) 3D cryo-EM map of the three-dimensional C9$^{CARD}$ filament at 3.3 Å resolution with an axial rise of 9.27 Å, pitch 49.04 Å, 5.3 units/turn. (**C**) Representative cryo-EM micrograph of C9$^{CARD}$ (WT) filaments used for structure determination. (**D**) Schematic representation of the hexagonal assembly of the C9$^{CARD}$ domain within the filament structure showing three asymmetric interfaces with opposing surfaces designated as 'a' and 'b'. (**E**) GS-FSC between half maps and map *vs* model at 0.143 and 0.5 threshold respectively.

MDEADRRLLRRCRLRLVEEL$^{20}$ QVDQLWDALLSRELFRPHMI$^{40}$

EDIQRAGSGSRRDQARQLII$^{60}$ DLETRGSQALPLFISCLEDT$^{80}$

GQDMLASFLRTNRQAAKLS$^{99}$

**Supplementary Figure 8**. Sequence of the C9$^{CARD}$ residues 1-99 with the negatively (D, E) and positively charged (R, K) residues shown in red and blue, respectively. The H38 residue has been highlighted. The overall net charge at pH 7 is expected to be zero due to the equal numbers of charged residue types – 16 negative (9 D, 7 E) and 16 positive (15 R, 1 K). The construct included a Ser-Gly overhang before Met1.

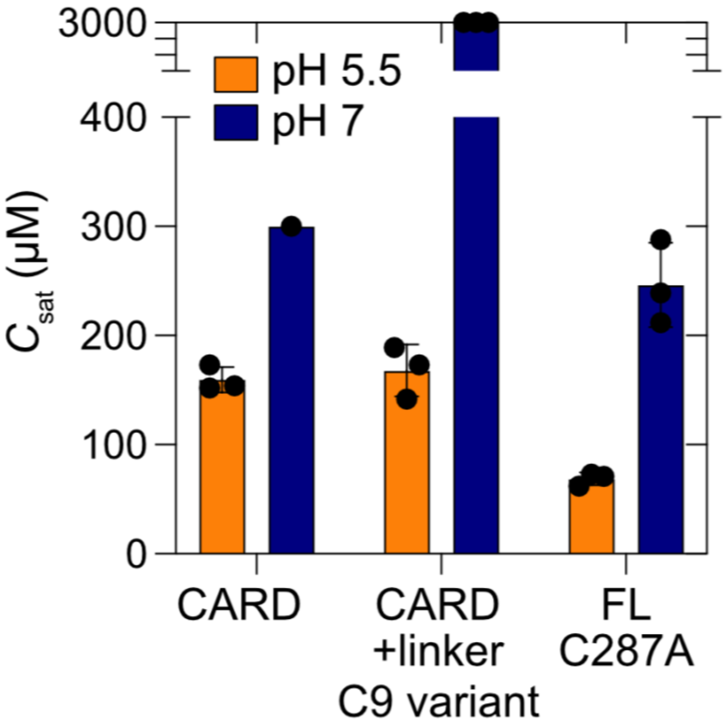


**Supplementary Figure 9.** Solubility of different C9 variants at two pH values. Measurements reflect three independent experiments for the $C9^{CARD}$, the $C9^{CARD+linker}$, or full-length C9 with the C287A mutation. The solution pH was either 5.5 (orange) or 7.0 (blue) and the experiment was performed at 25 °C.

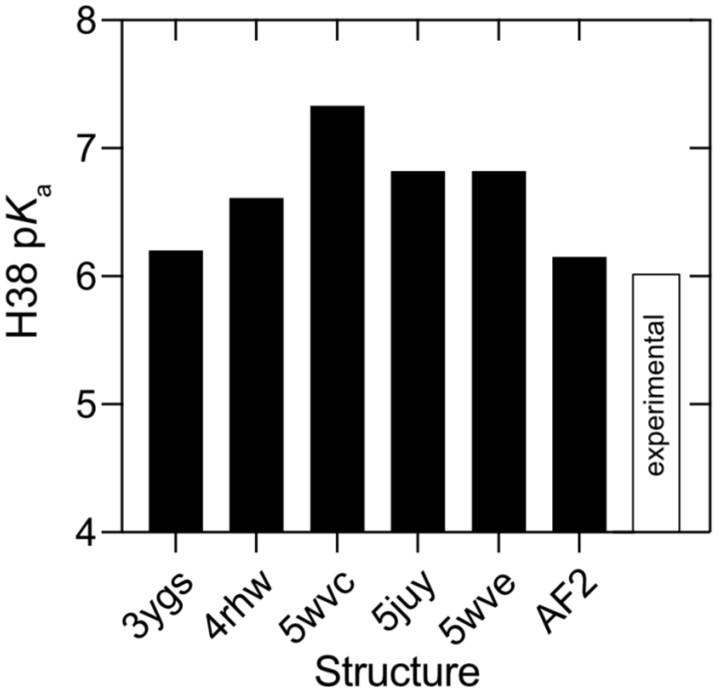


**Supplementary Figure 10. Structure-based H38 p$K_a$ prediction in the C9$^{CARD}$.** Structure-based prediction of the p$K_a$ value of H38 in the C9$^{CARD}$, shown as a function of the input structure (black, filled). AF2 corresponds to C9 model deposited the AlphaFold Protein Structure Database. The NMR-derived, experimentally determined p$K_a$ value for H38 is shown in white (via Figure 3).

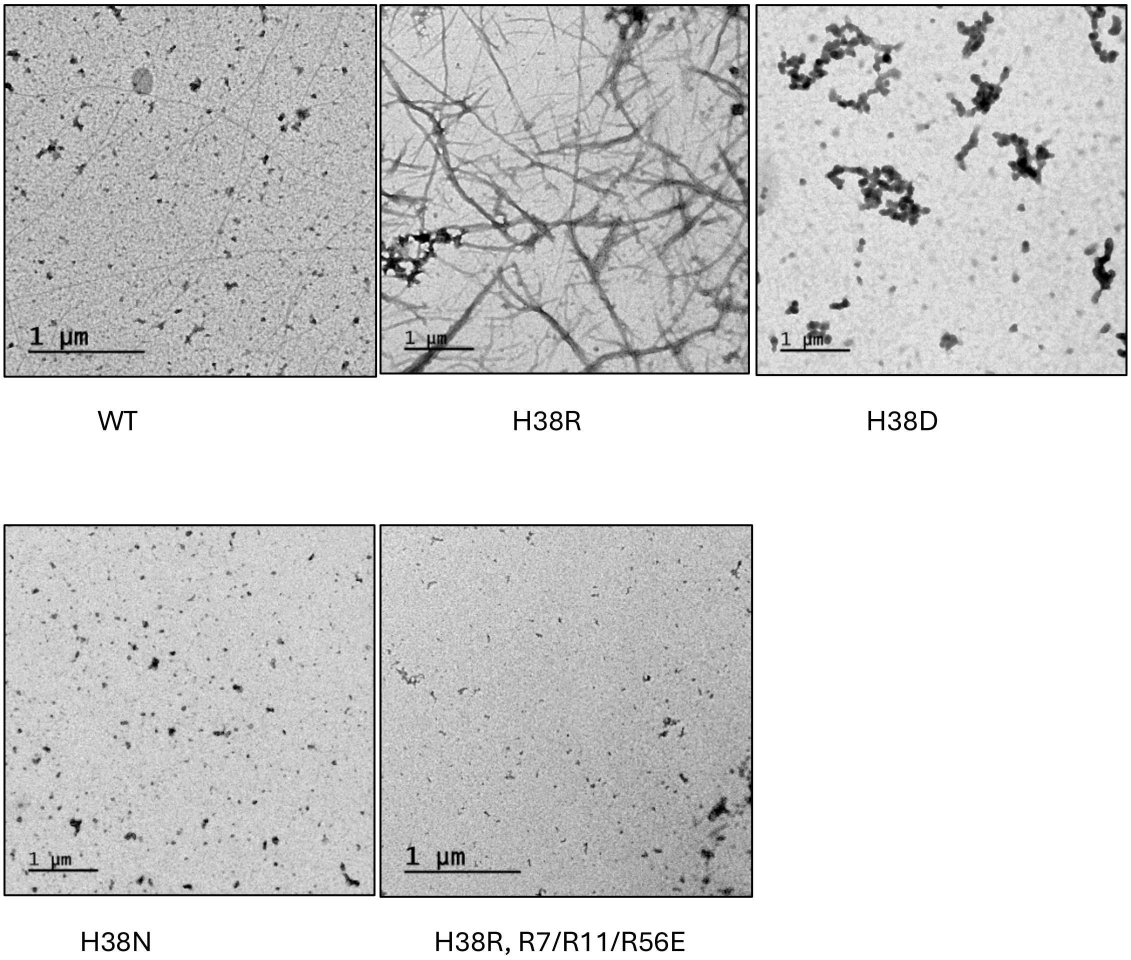


**Supplementary Figure 11**. Representative negative-stain micrographs of C9$^{CARD}$ variants in pH 5.5 buffer obtained after 40 minutes of centrifugation. Filaments are readily observed for the wild-type (WT) and H38R C9$^{CARD}$, whereas amorphous aggregates or no aggregates are observed for the H38D, H38N, and R7/R11/R56E + H38R variants under these conditions.

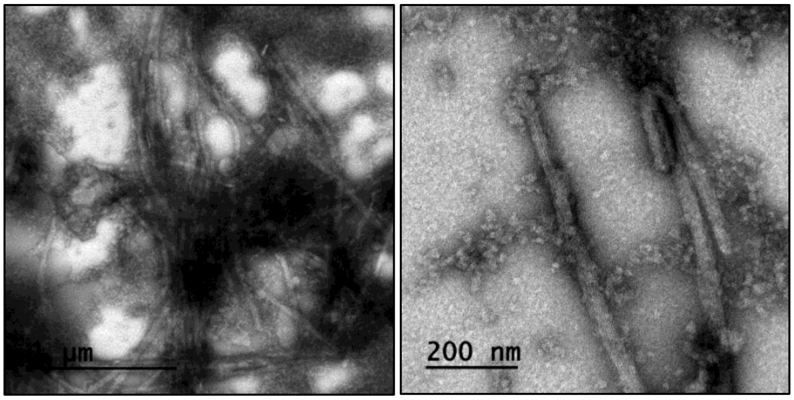


**Supplementary Figure 12**. Representative negative-stain EM micrographs of the H38D C9$^{CARD}$ variant at pH 5.5 obtained after 80 minutes of centrifugation where some filaments can now be observed.

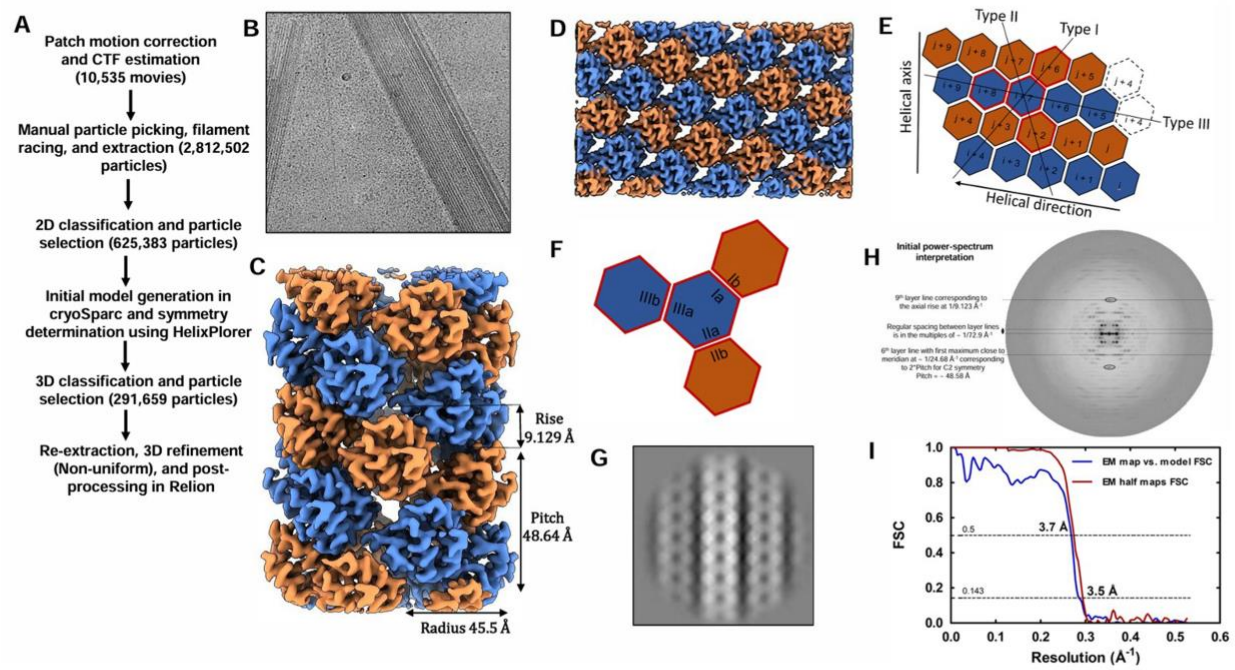


**Supplementary Figure 13. Structure determination of H38R $C9^{CARD}$ filament** (**A**) Data processing pipeline for 3D reconstruction using CryoSparc and Relion. (**B**) Representative cryo-EM micrograph of $C9^{CARD}$ (H38R) filaments used for structure determination. (**C**) 3D cryo-EM map of the three-dimensional $C9^{CARD}$ filament at 3.5 Å resolution. (**D-F**) Schematic representation of the 2D lattice and hexagonal assembly of the $C9^{CARD}$ domain within the filament structure showing three asymmetric interfaces with opposing surfaces designated as ‘a’ and ‘b’. (**G-H**) 2D-class average and the corresponding power spectra used for initial symmetry parameter estimation with initial interpretation on left and the indexing right corresponding to the final symmetry with axial rise 9.1 Å, pitch 48.6 Å, and 5.2 units/turn. (**I**) GS-FSC between half maps and map *vs* model at 0.143 and 0.5 threshold respectively.

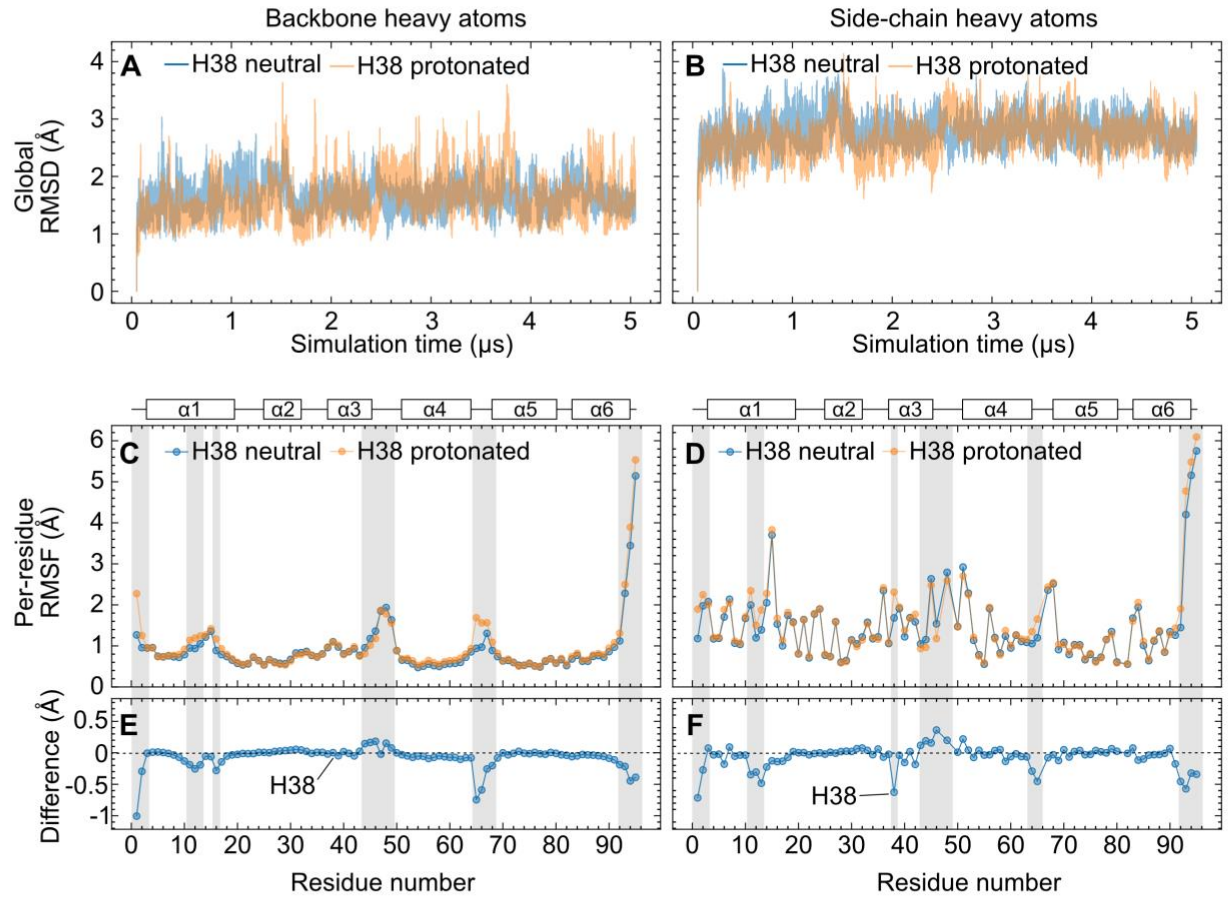


**Supplementary Figure 14. All-atom MD simulations of C9$^{CARD}$ with either uncharged or positively charged H38.** Simulations were performed using PDB 4rwh (chain E) that was energy-minimized and equilibrated at 300 K and 1 bar. The total duration of simulation time was 5.05 μs. The initial 50 ns were discarded, and the subsequent 5 μs were analyzed here. Neutral and protonated H38 correspond, respectively, to the singly protonated $N^{\varepsilon2}$–H τ tautomer and biprotonated $N^{\varepsilon2}$–H and $N^{\delta1}$–H state (cationic imidazolium). Root-mean-square deviation (RMSD) values of backbone (**A**) and side-chain (**B**) heavy atoms are shown as a function of the simulation time. In both protonation states of H38, the overall fold of the protein is stable with only local structural fluctuations. Per-residue root-mean-square fluctuation (RMSF) values are shown in (**C**) and (**D**) for the backbone and side-chain heavy atoms, respectively. The differences, H38 neutral – H38 protonated, for the backbone and side-chain heavy atoms are shown in (**E**) and (**F**), respectively, with negative values corresponding to higher RMSF values in the protonated state. Residues with significant changes to their RMSF values upon H38 protonation are indicated with grey boxes. This includes the side chain of H38 itself (**F**). The most significant changes to the backbone localize to the N-terminus (M1, D2), the C-terminal region of α3 and the α3-α4 loop (Q44, R45, A46, S48), and the α4-α5 loop (R65, G66, S67, Q68). The network of interactions involving H38-M39-R65/G66-M1 is sensitive to H38 protonation. Regions of secondary structure are indicated above panels **C** and **D** for clarity.

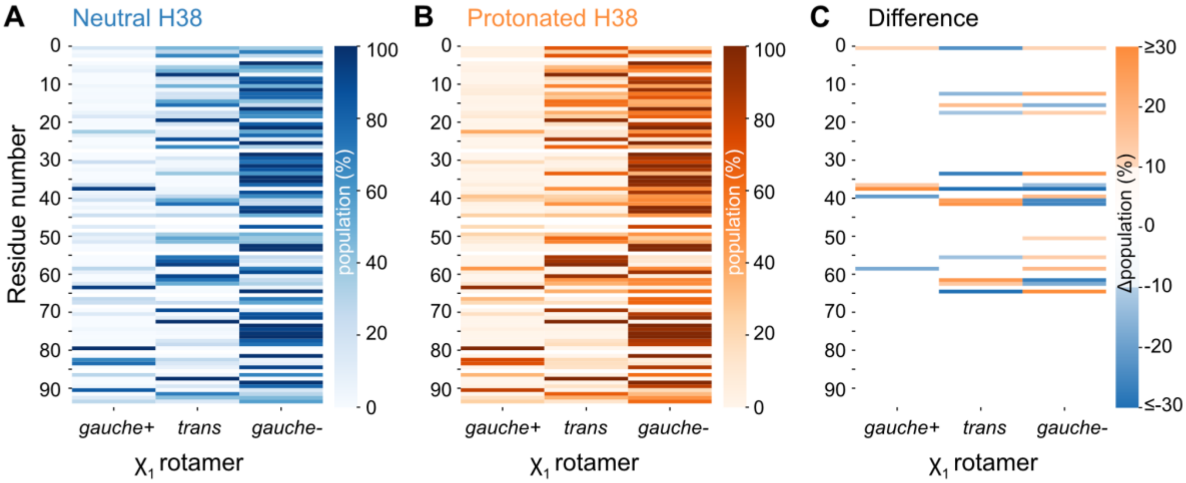


**Supplementary Figure 15. Protonation of H38 leads to a redistribution of side-chain $\chi_1$ torsion angles.** $\chi_1$ rotamer populations (*gauche*+, *trans*, *gauche*-) derived from the 5-μs MD simulations of the H38-neutral (**A**) or H38-protonated (**B**) states of the C9$^{CARD}$. Residues with empty data points correspond to Ala and Gly residues. (**C**) Population difference plot (protonated minus neutral), where orange (blue) values are positive (negative) and correspond to an enriched (depleted) rotamer in the H38-protonated state.

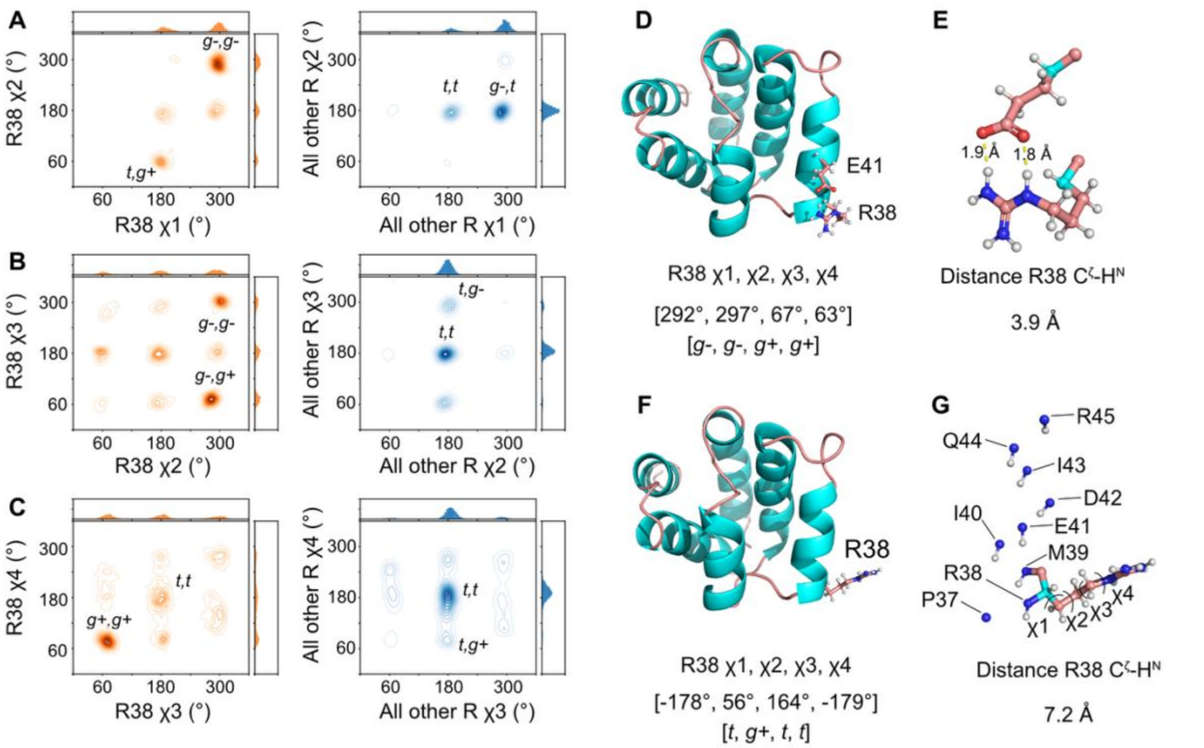


**Supplementary Figure 16. All-atom MD simulations of C9$^{CARD}$ with the H38R mutation.** Simulations were performed using PDB 4rwh (chain E) that was energy-minimized and equilibrated at 300 K and 1 bar. The total duration of simulation time was 5.05 μs. The initial 50 ns were discarded, and the subsequent 5 μs were analyzed here. The H38R mutation was introduced in PyMOL using the Mutagenesis tool. (**A**) (left) 2D histogram showing $\chi_1$ and $\chi_2$ rotamers of R38 throughout the simulation (orange). (right) The same plot for all other 15 Arg residues in C9$^{CARD}$ (blue). The two most populated $\chi_1$,$\chi_2$ states are labeled. The same plots are shown in (**B**) and (**C**), except for ($\chi_2$,$\chi_3$) and ($\chi_3$,$\chi_4$), respectively, noting that $\chi_4$ is non-rotameric but the closest states were assigned for consistency. (**D**) The most abundant $\chi_{1-4}$ angles produce the following conformation of R38 in which it is oriented in close proximity to E41, located *i*+3 in the helix, and forms a salt bridge. The distance between the R38 guanidinium group (R38 C$^{\zeta}$) and the helix dipole (R38 H$^{N}$) is 3.9 Å. (**F**) The same as (D) except for the second most abundant $\chi_{1-4}$ angles, leading to a conformation in which R38 is oriented toward solvent. (**G**) The distance between the R38 guanidinium group (R38 C$^{\zeta}$) and the helix dipole (R38 H$^{N}$) is 7.2 Å. The N-H groups in the helix are shown in spheres and the $\chi_{1-4}$ angles are annotated.

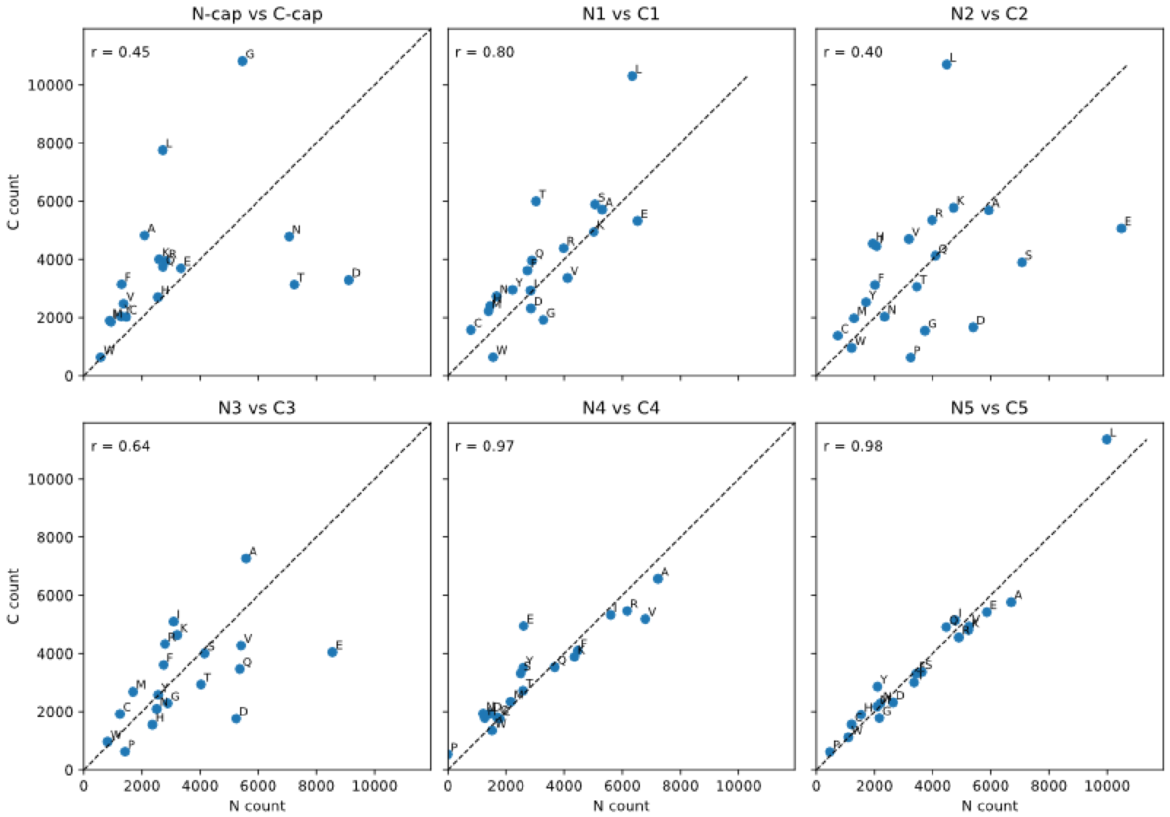


**Supplementary Figure 17. Correlation of helical residue types by helical position.** Shown here are the observed count of each residue type at the indicated helix residue position for N- (x-axis) vs. C-terminal (y-axis) positions. A minimum helix length of 12 residues was imposed on the AF2 ECOD helical residue atlas. These data are replicated in Figure 6 for the N1/C1, N2/C2, N3/C3, and N5/C5 positions. Note that N5/C5 here were defined as the helix middle. The Pearson correlation coefficient is indicated in the upper left corner, and the dashed line indicates y = x. Note that the C1 position does not contain P by definition, and therefore this point does not exist on the correlation plot. (It is indicated with an asterisk * in Figure 6). Also absent from the N3/C3 plot is L which is located at (7455, 13044).

### Supplementary Appendix

Apoptosis, or programmed cell death, is an intricate biochemical signaling cascade that eliminates damaged cells and aids in the regulation of cellular differentiation and homeostasis[1–3]. Dysregulated apoptosis plays a central role in unregulated inflammatory responses, neurodegenerative diseases, and cancer development[4–7]. Caspase-9 (C9), a cysteine-aspartic protease[8,9], plays a key role as an initiator of the intrinsic apoptotic pathway[10,11] and regulates cell death processes by cleaving and activating downstream effector caspases, such as C3 and C7 [12–14]. Under basal conditions, C9 is present as an inactive monomer and requires clustering and activation by the apoptosome[10,15,16], a heptameric scaffold comprising Apaf-1, ATP, and cytochrome C. While the role of C9 in activating effector caspases is well established, it is increasingly clear that C9 cleaves hundreds of substrates with diverse implications[17,18].

The CARD-containing protease caspase-9 (C9) plays a key role as an initiator of the intrinsic apoptotic pathway[10,11] and regulates cell death processes by cleaving and activating downstream effector caspases, such as C3 and C7 [12–14]. Under basal conditions, C9 is present as an inactive monomer and requires clustering and activation by the apoptosome[10,15,16], a heptameric scaffold comprising Apaf-1, ATP, and cytochrome C. While the role of C9 in activating effector caspases is well established, it is increasingly clear that C9 cleaves hundreds of substrates with diverse implications[17,18]. Self-assembly in C9 has mainly focused on its protease domain, for which substrate-driven, proximity-induced dimerization on the apoptosome platform regulates enzyme activity[19,20]. Protein engineering approaches demonstrated control over the self-assembly and activation of the C9 protease domain via constitutive protease-domain dimerization, recruitment to synthetic oligomeric platforms, or by fusion to a chemically-inducible dimerization domain as a safety switch in adoptive T-cell therapy[21,22,23 24]. Moreover, the activation or catalytic activity of C9 can be inhibited allosterically by post-translational modifications that are distal from the active site [25–30], as well as by zinc and peptide binding to other allosteric sites on the protease domain[31,32]. More generally, allostery in caspase protease domains appears to be widespread [33–40], including in C9 where it displays half-of-sites reactivity [19,20]. The $C9^{CARD}$ mediates recruitment to the apoptosome platform via CARD-CARD interactions with $Apaf\text{-}1^{CARD}$ that involve three different interfaces [41–43]. The CARD-CARD complex forms a helical oligomer or disk[15,41,42,44–46], and the same helical-disk architecture involving C9$^{CARD}$ and Apaf-1$^{CARD}$ has been observed in cryo-electron microscopy (EM) structures of the apoptosome:caspase-9 complex[43,47].

At the molecular level, C9 is a 416-residue protein that is comprised of a caspase activation and recruitment domain (CARD, residues 1-96), an interdomain linker, and a protease domain (residues 139-416). Activation of the protease domain involves substrate-driven, proximity-induced dimerization on the apoptosome platform[19,20] . Protein engineering approaches have further demonstrated that C9 can be activated via recruitment to synthetic platforms or by constitutive dimerization of the protease domain[21,22,23] . The dimerization-dependent activity of C9 was leveraged as a safety switch in adoptive T-cell therapy, whereby the expression of a chimeric protein comprising the protease domain of C9 and a chemically-inducible dimerization domain enables targeted and rational control over C9 activation and cell death[24]. Moreover, the activation or catalytic activity of C9 can be inhibited allosterically by post-translational modifications that are distal from the active site [25–30], as well as by zinc and peptide binding to other allosteric sites on the protease domain[31,32]. More generally, allostery in caspase protease domains appears to be widespread [33–40], including in C9 where it displays half-of-sites reactivity [19,20], which is an extreme form of negative cooperativity.

The N-terminal CARD domain of C9 belongs to the death-domain fold (DDF) superfamily, which fold into conserved six-helix bundles and mediate protein-protein interactions[48–50]. The DDF superfamily comprises four sub-families that include the CARD, death domain (DD), death effector domain (DED), and pyrin domain (PYD). DDFs primarily make homotypic interactions within sub-families (CARD-CARD, PYD-PYD, etc.) with remarkable specificity[49]. The C9 CARD (C9$^{CARD}$) mediates recruitment to the apoptosome platform via CARD-CARD interactions with Apaf-1$^{CARD}$. The CARD-CARD complex forms a helical oligomer or disk [15,41,42,44–46], and the same helical-disk architecture involving C9$^{CARD}$ and Apaf-1$^{CARD}$ has been observed in cryo-electron microscopy (EM) structures of the apoptosome:caspase-9 complex[43,47]. In full-length C9, the CARD appears to tumble independently of the protease domain[20], although transient inter-domain interactions may influence substrate recruitment[51]. CARDs often use multiple interfaces to interact with themselves and others, including in C9 where the C9$^{CARD}$:Apaf-1$^{CARD}$ complex involves three different interfaces[41–43]. The locally asymmetric binding interfaces among CARD complexes can lead to complex modes of homo- and hetero-oligomerization, such as polymerization into helical filaments [52–61] and the formation of biomolecular condensates via liquid-liquid phase separation[62]. For caspases, the formation of supramolecular complexes via DDF self-assembly dramatically increases the local concentration of their protease domains, leading to rapid activation. For example, filament formation was shown to play a role in the activation of the CARD-containing C1[53] as well as the tandem DED-containing C8[63,64].
